# Frustration Landscapes of Broadly Neutralizing SARS-CoV-2 Spike Antibodies Targeting Conserved Epitopes Reveal Energetic Logic of Escape-Proof and Escape-Prone Mechanisms

**DOI:** 10.64898/2026.04.02.716254

**Authors:** Mohammed Alshahrani, Will Gatlin, Max Ludwick, Lucas Turano, Brandon Foley, Gennady Verkhivker

## Abstract

The continued evolution of SARS-CoV-2 has enabled escape from most monoclonal antibodies, yet a subset of broadly neutralizing antibodies targeting three newly identified super-conserved RBD epitopes—SCORE-A, SCORE-B, and SCORE-C—retains remarkable activity against even the most recent JN.1-derived sublineages. Here we employed an integrated computational framework combining conformational dynamics, mutational scanning, MM-GBSA binding energetics, and frustration profiling to dissect the molecular mechanisms by which XGI antibodies achieve broad neutralization and resistance to immune escape. Structural analysis revealed that all three SCORE epitopes share a common architecture: a highly conserved, minimally frustrated core that provides stable anchoring, flanked by peripheral regions that accommodate antibody-specific variations. Conformational dynamics showed that SCORE-A antibodies (XGI-183) rigidify the lateral epitope while leaving the RBM partially mobile; SCORE-B antibodies (XGI-198, XGI-203) clamp the RBM apex, directly blocking ACE2; and SCORE-C antibodies (XGI-171) allosterically loosen the RBM loop, impairing receptor engagement indirectly. Mutational scanning identified a hierarchical hotspot organization where primary hotspots (e.g., K356, T500, Y380, T385) are evolutionarily constrained and minimally frustrated, while secondary hotspots (e.g., V503, Y508, S383) are neutrally frustrated and represent the principal sites of immune-driven mutations. MM-GBSA decomposition revealed that van der Waals-driven hydrophobic packing dominates binding, with electrostatic interactions providing auxiliary stabilization. Critically, frustration analysis demonstrated that immune escape hotspots reside precisely in zones of neutral frustration—“energetic playgrounds” that permit mutational exploration without destabilizing the RBD—while minimally frustrated cores are evolutionarily locked. The comparative analysis of conformational versus mutational frustration distributions revealed a unifying principle: aligned neutral frustration yields permissive, escape-prone interfaces; decoupling enables targeting of constrained cores; and convergence of minimal frustration in both distributions creates invulnerable interfaces. These findings establish that broad neutralization arises not from ultra-high-affinity anchors but from strategic energy distribution across rigid, evolutionarily informed interfaces, providing a roadmap for designing next-generation therapeutics that target the invulnerable cores of viral surface proteins.

## 1. Introduction

The emergence of severe acute respiratory syndrome coronavirus 2 (SARS-CoV-2) has prompted an intensive global research effort to understand its molecular architecture, mechanisms of host cell entry, and the immune responses it elicits. Central to these investigations is the Spike (S) glycoprotein, a trimeric surface structure that mediates viral entry and serves as the primary target of neutralizing antibodies [1–15]. The S protein exhibits remarkable conformational flexibility, transitioning through multiple functional states—from receptor engagement to membrane fusion—while simultaneously evading immune surveillance [10–15]. Comprising two distinct subunits, S1 and S2, the S protein includes within S1 the N-terminal domain (NTD), the receptor-binding domain (RBD), and two conserved subdomains (SD1, SD2) that stabilize the prefusion conformation [16–18]. The RBD, in particular, plays a critical role in binding the angiotensin-converting enzyme 2 (ACE2) receptor, making it a focal point for neutralizing antibody responses. Extensive cryo-electron microscopy (cryo-EM) and X-ray structures of SARS-CoV-2 S protein variants of concern (VOCs) in various functional states, along with their interactions with antibodies, underscore how VOCs can induce structural changes in the dynamic equilibrium of the S protein [19–25]. These findings underscore the balance between structural stability, immune evasion, and receptor binding that shapes the evolutionary trajectory of SARS-CoV-2 and its variants.

The evolution of SARS-CoV-2 variants such as XBB.1 and XBB.1.5 have garnered significant attention due to their enhanced growth advantages, transmissibility, and immune evasion capabilities. These subvariants represented critical milestones in the virus’s evolutionary trajectory, highlighting its ability to adapt through mutations that optimize receptor binding while evading immune defenses [26,27]. XBB.1.5, a descendant of the BA.2 lineage, arose through recombination events that introduced key mutations in the RBD of the S protein. These mutations significantly enhance its binding affinity for the ACE2 receptor, making it more infectious than earlier Omicron strains. The continued evolution of SARS-CoV-2 within the Omicron lineage and its descendants such as XBB.1, XBB.1.5, and more recent subvariants (e.g., JN.1, KP.2, KP.3), highlights not only the virus’s extraordinary adaptability but also the emergence of convergent evolutionary hotspots—specific residues that are repeatedly and independently mutated across geographically and temporally distinct lineages [28,29]. The recurrence of identical or functionally similar mutations F456L, R346T, L455F, and K444T across unrelated variants signals a narrowing evolutionary landscape, where SARS-CoV-2 is increasingly optimizing a limited set of high-impact changes to balance transmissibility, immune evasion, and structural stability [26–29]. The JN.1-derived subvariants KP.2 and KP.3 have independently acquired a constellation of key S mutations—including R346T, F456L, Q493E, and V1104L—that collectively enhance both transmissibility and the ability to evade neutralizing antibodies. Additional offshoots of JN.1, such as LB.1 and KP.2.3, further illustrate this dynamic evolutionary pattern : they share hallmark mutations R346T and F456L, while also introducing distinct changes S:S31- and S:Q183H in LB.1, or S:H146Q in KP.2.3 [26–29]. Notably, the F456L substitution has emerged as a pivotal driver of immune escape, with KP.3 recognized as one of the most antibody-evasive sublineages within the JN.1 clade [29]. This rapid diversification underscores extraordinary capacity of the virus to adapt under immune pressure, fine-tuning mutations to balance evasion with essential functional requirements such as receptor binding. Recent cryo-EM structural analyses of RBD complexes from JN.1, KP.2, and KP.3 reveal that the F456L mutation synergizes with Q493E to strengthen ACE2 binding providing a structural basis for KP.3 heightened infectivity and immune resistance [30,31].

The recombinant variant XEC, which originated from KP.3, has recently drawn attention due to two additional mutations in the NTD F59S and T22N [32,33]. XEC displays markedly higher infectivity than its parental KP.3 lineage and exhibits enhanced resistance to neutralizing immune responses. Separately, the KP.3.1.1 subvariant emerged from KP.3 through a deletion at position S31 in the NTD. Structural and binding analyses of the KP.3.1.1 RBD in complex with ACE2 revealed a critical epistatic interaction between two key RBD mutations F456L and Q493E [33]. In recent months, the JN.1-derived LP.8.1 and LP.8.1.1 sublineages have now largely displaced XEC across Europe and North America. Concurrently, LF.7 and its descendant LF.7.2.1 gained prominence while MC.10.1 experienced a sharp, albeit brief, surge in global prevalence [34,35]. LF.7 carries seven additional Spike mutations relative to JN.1: four in the NTD : T22N, S31P, K182R, and R190S—and three in the RBD - R346T, K444R, and F456L Its successor, LF.7.2.1 variant acquired an additional A475V substitution in the RBD [35] and quickly outcompeted LF.7 across many geographical regions. More recent variants NB.1.8.1 and the recombinant XFG showed a pronounced growth advantage over earlier JN.1 sublineages. NB.1.8.1, emerging from the recombinant lineage XDV, harbors seven S mutations beyond those in JN.1, namely T22N, F59S, and G184S in the NTD, and A435S, F456L, K478I, and Q493E in the RBD [36,37]. XFG—a recombinant offspring of LF.7 and LP.8.1.2 currently under active monitoring—carries four additional mutations beyond LF.7: H445R, N487D, and Q493E in the RBD, and T572I in the SD1 [36,37]. As of March 2026, the recombinant lineage XFG (’Stratus’) which a chimeric recombinant of the LF.7 and LP.8.1.2 lineages and its primary sub-lineage XFG.2.5.1 [37,38] became the dominant global drivers. BA.3.2 represents a new lineage of SARS-CoV-2, genetically distinct from the JN.1 lineages (including LP.8.1 and XFG) is drawing increased attention from global public health officials as it spreads across multiple continents and demonstrates a notable ability to evade immune defenses. Relative to LP.8.1, BA.3.2 has 20 receptor-binding domain differences and 35 N-terminal domain differences and cryo-EM structures of the BA.3.2.1 S protein and its ACE2-bound complexes revealed distinctive structural features that mechanistically rationalize immune escape coupled to attenuated replication fitness [39]. Together, these variants illustrate that the evolutionary trajectory of SARS-CoV-2 has transitioned from discrete point mutations toward complex multilineage recombination events, where co-infection facilitates the rapid ‘shuffling’ of genomic segments enhancing transmissibility and immune escape.

The fundamental aspect of the immune response to SARS-CoV-2 is the production of antibodies that target various regions of the S protein, which plays a central role in viral entry into host cells. The humoral immune response to SARS-CoV-2 relies heavily on antibodies targeting the RBD. The Barnes classification divides SARS-CoV-2 RBD-targeting antibodies into four main structural classes based on their epitopes and binding to “up” or “down” conformations has been widely used to understand neutralization mechanisms, epitope mapping, and viral escape [40]. In this classification, class 1 recognizes “up” RBD conformations, overlapping with the ACE2-binding motif, class 2 bind a “down” RBD conformations, also targeting the ACE2-binding region; class 3 engages the side of the RBD, away from the receptor-binding motif; class 4 targets cryptic or conserved sites on the RBD underside, accessible only in the “up” conformation. Class 1 and 2 antibodies are the most potent and frequently elicited, targeting overlapping regions of the receptor-binding site (RBS) where ACE2 binds [40,41]. However, these antibodies exhibit limited cross-reactivity with related coronaviruses and readily escaped by SARS-CoV-2 variants. Class 3 and 4 antibodies are less potent and less frequently elicited in humans but target relatively more conserved RBD regions and exhibit cross-reactivity with VOCs [40,41].

High-throughput yeast display and deep mutational scanning (DMS) have enabled systematic mapping of antibody epitopes and escape mutations, leading to comprehensive classification of neutralizing antibodies into six major epitope groups (A–F) based on their binding footprints [42,43]. By integrating high-throughput DMS datasets, they engineered pseudoviruses using predicted escape mutations as selective filters to identify antibodies resilient to viral evolution [44]. A retrospective analysis of monoclonal antibodies originally derived from wildtype (WT) SARS-CoV-2 infection demonstrated the power of this approach as it boosted the likelihood of identifying antibodies effective against the XBB.1.5 variant from a mere 1% to 40% [45]. The six-group (A-F) classification, derived from large-scale DMS profiling captured nuanced differences in escape patterns where groups A-D correspond to Barnes classes 1 and 2, subdivided based on sensitivity to specific mutations where group E maps to Barnes class 3, targeting the RBD side with relative resilience to RBM mutations while group F corresponds to Barnes class 4, targeting conserved cryptic epitopes and demonstrating broad neutralization. Importantly, for our study, in the classification system developed by Cao and colleagues, class 4 antibodies (targeting the conserved “inner face” of the RBD) are subdivided into groups F1, F2, and F3 based on their specific epitopes, neutralization mechanisms, and susceptibility to mutations [42–45].

This functional subdivision proved critical for understanding how different antibodies within the same structural class exhibit distinct escape profiles, and for identifying antibodies with exceptional neutralization breadth. Additional classification systems have emerged, including the RBD-1 to RBD-7 framework, which provides a more granular breakdown while retaining the same hierarchy: RBD-1/2 correspond to Barnes class 1/2 and Cao groups A-D; RBD-3/4/5 correspond to Barnes class 3 and Cao group E; RBD-6/7 correspond to Barnes class 4 and Cao group F [46]. A granular structure-based classification exploited quantitatively defined contacting amino acid residues on the RBD as well as a clustering analysis. These analyses reveal common characteristics of some 23 frequently contacted ES and the structural nature of the surfaces of the RBD that interact with antibodies [47]. A systematic, large-scale analysis of SARS-CoV-2 antibody recognition^48^ used classification of all RBD-targeting antibodies into four broadly defined epitope groups based on residue overlap [47] where groups I and II target the highly variable receptor-binding site (RBS) and Groups III and IV engage more conserved, non-RBS regions. Collectively, this comprehensive analysis showed that antibodies cover 99% of the RBD surface, demonstrating that virtually every solvent-accessible residue can be targeted by mammalian antibodies pointing to resilience of the humoral immune response [48]. Several broadly neutralizing antibodies have been identified that maintain efficacy across diverse variants, revealing distinct mechanistic strategies. In particular, class 1 antibodies such as BD55-1205 [45], VIR-7229 [49], 19-77 [50], ZCP3B4 [51], and ZCP4C9 [51] achieve broad neutralization through different mechanisms. While most of clinically approved monoclonal antibodies have been rendered largely ineffective by the antigenic drift of Omicron sublineages to the evolving JN.1 subvariants, class 4 (group F3) antibodies pemivibart (VYD222) [52] and SA55 (BD55-5514) [53] showed remarkable neutralization breath against newest JN.1 descendant variants and target a similar RBD epitope and demonstrate broad neutralization against SARS-CoV-2 variants.

The continued emergence of escape variants underscored the importance of identifying and characterizing antibodies targeting truly conserved epitopes—sites where mutational escape imposes prohibitive fitness costs. A significant advance in this direction comes from the recent work of Cao and colleagues who systematically characterized orphan broadly RBD-binding antibodies and identified three conserved RBD epitopes that remain accessible throughout SARS-CoV-2 evolution [54]. Three groups of non-overlapping conserved epitopes on RBD were identified suggesting three non-competing super-conserved RBD epitopes, SCORE-A, SCORE-B and SCORE-C. Through comprehensive structural and functional analyses, including multiple high-resolution crystal structures, the study revealed that these epitopes represent vulnerable sites on the RBD that have remained largely invariant despite extensive antigenic drift in surrounding regions. The antibodies targeting these conserved epitopes, designated as XGI antibodies, fall within class 4 and site V categories and demonstrate remarkable neutralization breadth across diverse SARS-CoV-2 variants, including the most recent JN.1-derived sublineages [54]. Site V is another newly characterized highly conserved and relatively “silent” region on the RBD targeted by a special class of broadly neutralizing antibodies that resist extreme antigenic drift across variants [55]. Key examples include CC25.4, CC25.17, and CC25.56 antibodies which demonstrate broad neutralization with less potency than group 1 antibodies but significantly greater escape resistance [55] A few previously reported human antibodies including S2H97 [56], COVOX-45 [57], 553-49 [58], and XMA09 [59] also target this site and exhibit remarkably broad neutralization.

The interplay between structural dynamics and binding energetics is central to understanding how antibodies achieve broad neutralization and how viruses evolve to escape them. Computer simulations have significantly advanced our understanding of the dynamics and functions of the S complexes with ACE2 and antibodies at the atomic level [60–62]. Conformational dynamics and allosteric interactions can be linked to binding of novel human antibodies [63,64] where antibody-induced associated changes in S dynamics can distinguish weak, moderate and strong neutralizing antibodies [65]. Recent computational and structural studies suggested a mechanism in which the pattern of specific escape mutants for ultrapotent antibodies may be driven by a complex balance between the impact of mutations on structural stability, binding strength, and long-range communications [66,67]. At the molecular level, the evolution of immune escape hotspots in SARS-CoV-2 is influenced by both random genetic drift and natural selection acting on variants that provide fitness advantages. The concept of “energetic frustration”—regions where proteins are not fully optimized for stability but retain functional plasticity—has emerged as a framework for understanding why certain positions repeatedly mutate under selective pressure [68–71].

Our recent computational studies examined ultrapotent and broadly neutralizing Class 1 SARS-CoV-2 neutralizing antibodies—BD55-1205, 19-77, ZCP4C9, and ZCP3B4 and class 4/1 ADG20 antibody to reveal a unifying biophysical principle that explains how these antibodies achieve both exceptional potency and remarkable resilience against viral evolution [72,73]. Using multi-pronged computational approach, we found these neutralizing antibodies bind via rigid, pre-configured interfaces that distribute binding energy across a broad epitope through numerous suboptimal, yet synergistic, interactions—many of which occur at sites of neutral local frustration.

Structural analysis of XGI antibody-RBD complexes revealed that the three conserved epitopes are situated in regions under strong functional constraints where mutations at these positions would likely disrupt essential structural or functional properties, imposing significant fitness penalties. However, a detailed understanding of dynamics and energetics underlying unique neutralizing capacity of these antibodies is lacking. Additionally, balance of structural stability and adaptability when targeting conserved epitopes remains an important driver of neutralization antibody efficiency and quantifying molecular determinants and specific hotspots of immune escape-resistant targets is of significant interest. In the current study, we address these questions from a unified biophysical perspective by integrating structure, dynamics, and energetics analyses to investigate how the conserved binding epitopes interact with class 4 and site V antibodies throughout SARS-CoV-2 evolution. The structural characterization of these epitopes provides a unique opportunity to understand the molecular basis of sustained vulnerability in an otherwise highly mutable domain.

We employ a multi-pronged computational approach that integrates conformational dynamics of antibody-RBD complexes, systematic mutational scanning to identify binding hotspots and predict escape mutations, MM-GBSA binding free energy calculations with residue-based decomposition and energy landscape-based frustration analysis to examine adaptive evolution. Our analysis focuses on several key questions: (a) what structural and energetic features enable XGI antibodies to maintain binding across diverse variants?; (b) how do the three conserved epitopes differ in their accessibility, dynamics, and mutational vulnerability? (c) what are the predicted escape pathways for these antibodies, and what fitness costs might such mutations incur? (d) how do XGI antibodies compare to previously characterized broadly neutralizing antibodies in terms of binding mechanisms and resilience? Our study aims to identify vulnerabilities that may remain accessible despite continued viral adaptation, demonstrating that the evolutionary trade-offs SARS-CoV-2 faces—balancing immune evasion against the need to maintain ACE2 binding—are often based on energetically neutral, suboptimal variations that are highly context-dependent even for antibodies targeting the same class of epitopes.

Our results show that the dominant pattern of neutral frustration in the evolutionary hotspots on the RBD may create a state of “energetically suboptimal, adaptable frustration” that limits access to potentially superior adaptive outcomes but generates mutational “hotspots” - where mutations repeatedly and predictably occur, leading to highly repeatable evolutionary trajectories. The emerging understanding of the immune evasion mechanisms suggests that at the molecular level, the evolution of immune escape hotspots in SARS-CoV-2 which is a complex process can be influenced by both random drift of mutations due to neutrally frustrated architecture and natural selection that acts on these variants, favoring convergent evolution for those positions that provide a recurrent fitness advantage

## 2. Results

### 2.1 Structural Characterization of the Conserved RBD Epitopes Targeted by XGI Antibodies

Our study analyzed the S-RBD in complex with four groups (A–D) of neutralizing antibodies, which exhibit diverse binding mechanisms and epitope specificities impacting neutralization potency and viral escape susceptibility. Group A antibodies, like CB6/LY-CoV016 bind exclusively to the ‘up’ RBD conformation [74]. LY-CoV016’s epitope spans residues 403–505, overlapping significantly with the ACE2 binding interface, blocking viral entry via charged residues, hydrophobic contacts, and salt bridges, notably K417 (Figure 1A, Table S1) [59,74]. Escape mutations primarily affect heavy-chain CDR-interacting residues: K417, D420, L455, F456, Y473, A475, N487, G504 [59].

**Figure 1.**
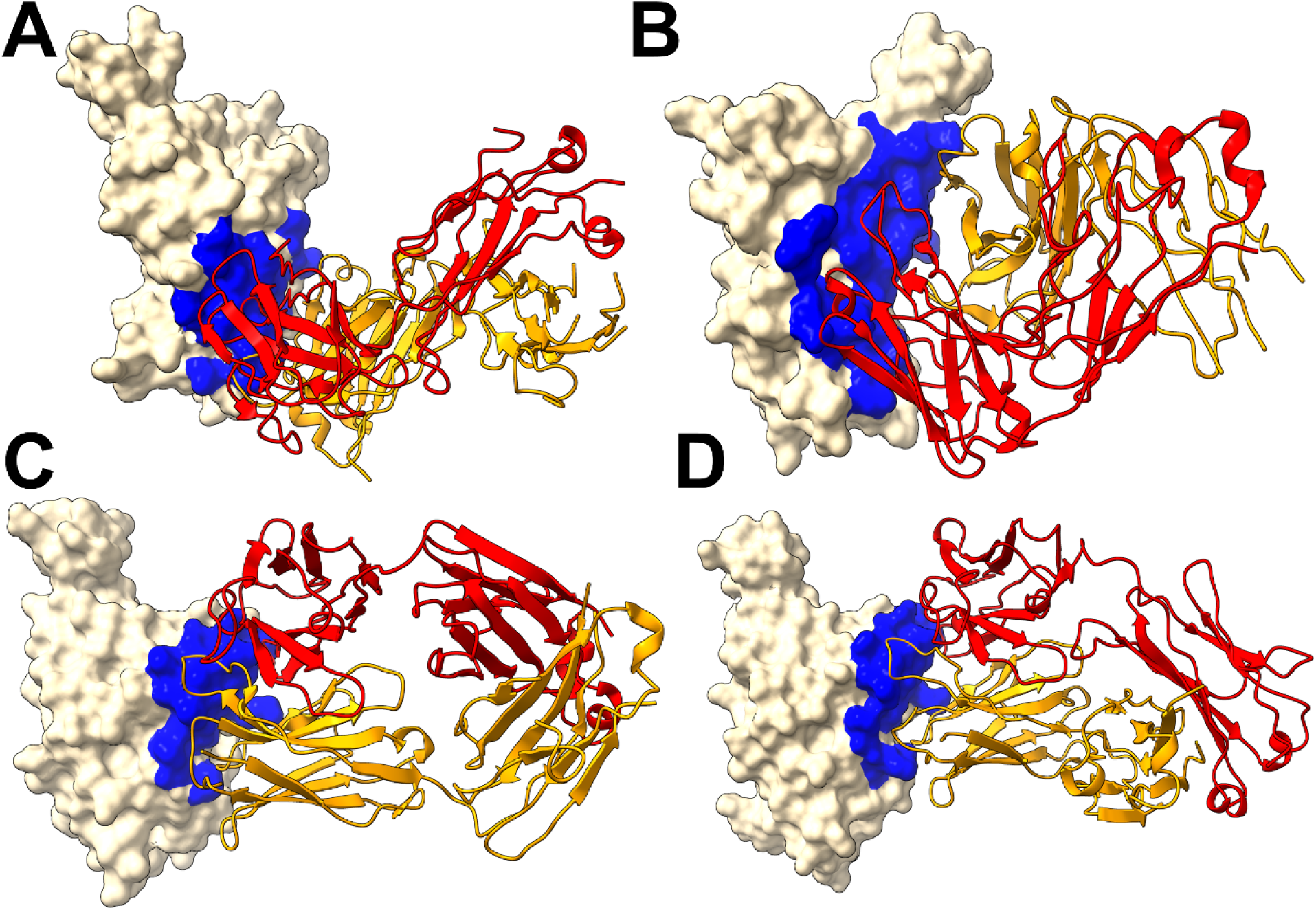
Structural organization of the RBD complexes for XGI antibodies. (A) The structure of XGI-171 with RBD (pdb id 9KZD). (B) The structure of class XGI-183 bound with RBD (pdb id 9KZE). (C) The structure of class XGI-188 bound with RBD (pdb id 9L05). (D) The structure of class XGI-203 bound with RBD (pdb id 9L07).). The RBD in wheat-colored surface. The epitope sites are highlighted in blue surface. The antibody heavy chain in orange ribbons, and the light chain in red ribbons.

We first performed structural analysis of four representative XGI antibodies in complexes with the Omicron EG.5.1 spike: XGI-183 (SCORE-A, PDB 9KZE), XGI-203 (SCORE-B, PDB 9L07), XGI-198 (SCORE-B, PDB 9L05), and XGI-171 (SCORE-C, PDB 9KZD) [54] (Figure 1). We follow the terminology and annotation from the original study that identified three non-overlapping super-conserved RBD epitopes, SCORE-A, SCORE-B and SCORE-C [54]. The structural analysis reveals a common architecture across all three epitopes: a highly conserved, minimally frustrated core that provides stable anchoring, flanked by peripheral regions that accommodate antibody-specific variations and determine neutralization potency and escape vulnerability. The three conserved RBD epitopes—SCORE-A, SCORE-B, and SCORE-C—are each recognized by distinct families of broadly neutralizing antibodies [54] XGI-171 targets the SCORE-C epitope characterized by cryptic inner face of the RBD accessible only when the RBD adopts the “up” conformation (Figure 1A).

The SCORE-A epitope is located on the lateral side of the RBD, adjacent to the N-terminal domain interface, and is targeted by a family of broadly reactive antibodies that includes XGI-183 [54] as well as S309 antibody [75–77], and SA58 (also known as BD55-5840) [43,78]. All three antibodies engage a central region of the RBD comprising residues 340–360 (α2-helix and the adjacent loop) and residues 457–471 (the β4-β5 hairpin and the loop preceding the α3-helix). (Figure 1B). The SCORE-B epitope (XGI-188, XGI-203) is at the tip of the RBD, partly overlapping the receptor-binding motif (RBM) and the ACE2-interacting surface. Antibodies XGI-188, XGI-203 targeting this region can directly block ACE2 binding, and their neutralization potency often correlates with the extent and geometry of epitope engagement (Figure 1C,D). Structural study showed that these three identified non-overlapping SCOREs correlated with the epitopes of S309 and SA58 for SCORE-A, SA55 for SCORE-B, and CR3022 and EY6A for SCORE-C, respectively [54]. We present a comparative analysis of binding epitopes for XGI antibodies alongside related antibodies (Figure 2). In XGI-183, the network of interactions is broad and extensive (Figure 2A, Supporting Information Table S1). Residues E340, V341, N343, A344, T345, T346, N354, R355, K356, R357, I358, S359, N360 form a dense network of hydrogen bonds and van der Waals contacts with the antibody heavy chain (CDR H2 and H3) and light chain (CDR L1 and L3).

**Figure 2.**
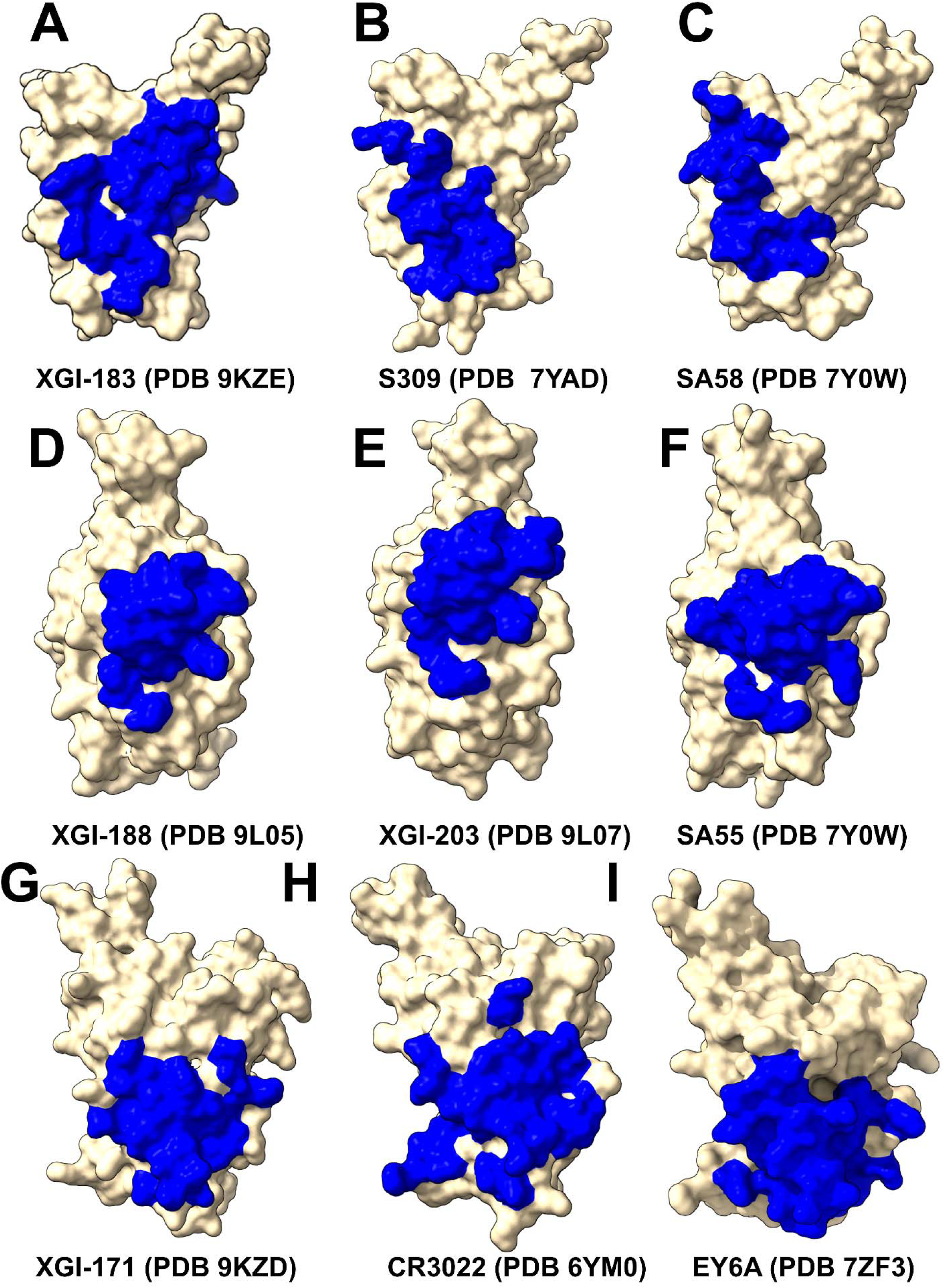
Structural Epitope Mapping of XGI Antibodies and SCORE-A,B,C Antibodies Binding to the RBD. (A-C) The binding epitopes for three representative SCORE-A antibodies - XGI-183 (PDB ID: 9KZE), S309 (PDB ID: 7YAD), SA58 (PDB ID: 7Y0W). (D-F) The binding epitopes for three representative SCORE-B antibodies - XGI-188 (PDB ID: 9L05), XGI-203 (PDB ID: 9L07), SA55 (PDB ID:7Y0W). (G-H) The binding epitopes for three representative SCORE-C antibodies - XGI-171 (PDB ID:9KZD), CR3022 (PDB ID: 6YM0), EY6A (PDB ID: 7ZF3). The RBD is depicted in wheat-colored surface, with the heavy chain (HC) of the antibody in orange and the light chain (LC) in red. The binding epitope residues are shown in blue surface. These panels reveal that while all three antibodies target a deeply conserved hydrophobic core within the RBD, there are appreciable differences in their contact footprints and residue-specific interactions.

XGI-183 exhibits a particularly extensive interaction with the 340–346 region. T345 forms multiple hydrogen bonds with light-chain residues N26, N27, and heavy-chain residues D38, G57, T58, D64, Y66, and W113; K356 contacts eight residues on the light chain (I28, G29, S36, K37, N38, V39, D57, N80) and is critical for binding (Figure 2A, Supporting Information Table S1). Another distinguishing feature of XGI-183 is its interaction with the 457–471 region. I468 and T470 make extensive contacts with the heavy chain (I468 with V55, N64, K65, H66, and light-chain W107, F113, D114, W116; T470 with K65, H66, K72, and light-chain F113). These residues are highly conserved and are not targeted by S309 (Figure 2B, Supporting Information Table S2) or SA58 (Figure 2C, Supporting Information Table S3) to the same extent.

S309 is unique among SCORE-A antibodies in that it interacts with the N343 glycan, which is conserved across sarbecoviruses. SA58 (BD55-5840) engages a slightly more “right-shifted” epitope compared to XGI-183 and S309, with contacts extending to residues L441, K444, N450, and R509 (Figure 2C, Supporting Information Table S3). Overall, XGI-183 relies on a broad protein-protein interface with a strong dependence on K356 while S309 achieves high potency through a combination of protein contacts and glycan-mediated stabilization, and SA58 balances broad binding with increased sensitivity to mutations at the periphery of its epitope. Structures of antibodies XGI-198 (PDB 9L05), XGI-203 (PDB 9L07) and SA55 (PDB 7Y0W) bound to RBD reveal shared and unique features of targeting the SCORE-B epitope residing closer to the tip of the RBD and partly overlapping with the ACE2-interacting surface (Figure 2D-F). These antibodies bind to a conserved core comprising residues 437–445 (β5-β6 loop) and 498–508 (RBM apex), all of which are essential for ACE2 binding (Figure 2D-F). Shared core interactions involve N437, N439, K440, K444, P445, R498, P499, T500, Y501, G502, V503, G504, and Y508. These residues are contacted by all three antibodies, primarily through the heavy chain (Figure 2D-F, Supporting Information Tables S4-S6). XGI-198 uniquely engages Q506 through multiple heavy-chain residues (H38, H57, H64, H66, H109, H112) and light-chain residue L114 (Figure 2D, Supporting Information Table S4). XGI-203 lacks strong engagement with Q506 and instead makes stronger contacts with residues A372, P373, F374, F375, S375, T376 (Figure 2E, Supporting Information Table S5). SA55 engages a broader set of residues, including R403, D405, V407, R408 on the conserved structural “mesa” adjacent to the ACE2 interface (Figure 2F, Supporting Information Table S6). XGI-171 that targets SCORE-C epitope engages a highly conserved hydrophobic core formed by residues Y369, F377, C379, Y380, G381, V382, S383, P384, T385, K386, D427, and D428 (Figure 2G, Supporting Information Table S7), particularly making extensive contacts with the 385–386 region. T385 interacts with seven heavy-chain residues (D38, I56, G57, T58, D64, Y66, W113) and K386 with five heavy-chain residues (D38, G108, S109, G110, Q111). Notably XGI-171 does not contact the 515–519 β7 strand nor the 412–414 loop, which are targeted by SCORE-C antibodies CR3022 (Figure 2H, Supporting Information Table S8), and EY6A (Figure 2I, Supporting Information Table S9).

Together, these structural analyses reveal that XGI antibodies achieve their remarkable breadth by anchoring to evolutionarily constrained, structurally stable cores, while their unique interfacial patterns—especially the extensive engagement of specific residues (K356 in XGI-183, Q506 in XGI-198, T385/K386 in XGI-171)—dictate their individual neutralization potency and escape vulnerability.

### 2.2. Conformational Dynamics of XGI Antibody–RBD Complexes Reveal Distinct Flexibility Signatures Linked to Escape Vulnerability

We investigated the conformational dynamics of the XGI antibody–RBD complexes using coarse-grained (CG-CABS) [79] and all-atom MD simulations. Root-mean-square fluctuations (RMSF) of Cα atoms provide a residue-level view of local flexibility, revealing how antibody binding modulates the intrinsic motions of the RBD. By comparing the RMSF profiles of XGI antibodies—XGI-171 (SCORE-C), XGI-183 (SCORE-A), XGI-203 (SCORE-B), and XGI-198 (SCORE-B)—and by placing them in the context of well-characterized antibodies from the same epitope classes (CR3022 and EY6A for SCORE-C; S309 for SCORE-A; SA55 for SCORE-B), we examine relationships between the dynamic signature of the RBD and the neutralization mechanism (Figure 3). XGI-183 binds the lateral side of the RBD, engaging two distinct clusters: the α2-helix (residues 340–360) and the β4-β5 hairpin (residues 457–471) (Figure 3A). Both regions are significantly stabilized, with RMSF values in the range of 0.2–0.6 Å for key residues such as K356, R357, I468, and T470. This rigidification provides a stable anchor. The RBM loop (470–490) shows intermediate flexibility, with peaks around 2.5–3 Å—less than in SCORE-C complexes but clearly higher than in SCORE-B. The α2-helix and β4-β5 hairpin are thus locked, while the RBM loop remains partially mobile. This dynamic pattern reflects a dual mechanism: the antibody directly stabilizes the lateral epitope and partially restricts the RBM but does not fully clamp the ACE2-binding surface (Figure 3A).

**Figure 3.**
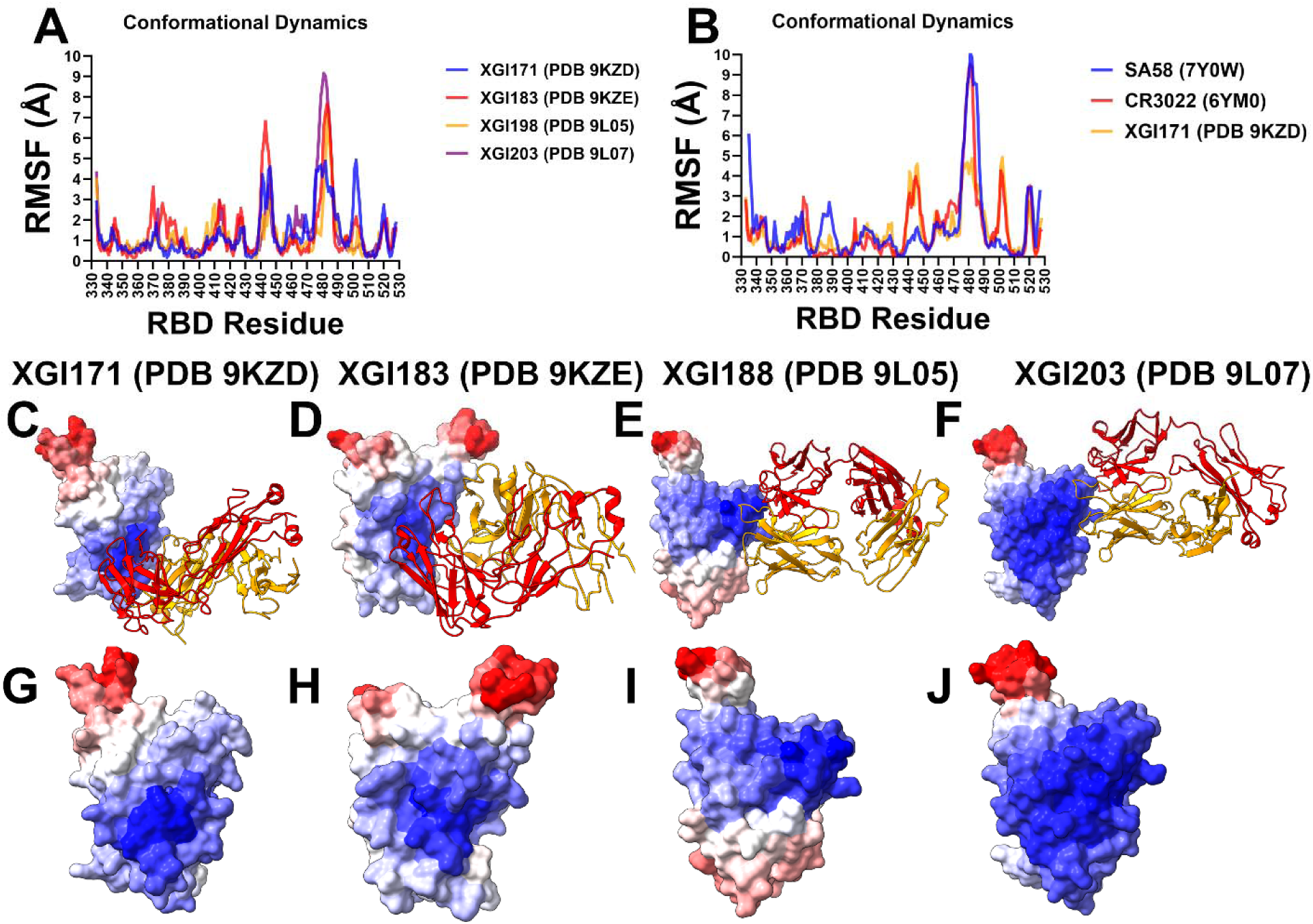
Conformational dynamics profiles obtained from CG-CABS simulations and atomistic reconstruction of the RBD-antibody complexes. (A) The RMSF profiles for the XGI antibodies : XGI-171 with EG.5.1 RBD (in blue lines), XGI-183 with EG.5.1 RBD (in red lines), XGI-188 with EG.5.1 RBD (in orange lines), and XGI-203 with EG.5.1 RBD (in purple lines). (B) The RMSF profiles for the SCORE-C antibodies : XGI-171 with EG.5.1 RBD (in orange lines), CR3022 with RBD with RBD (in red lines) and EY6A with RBD (in blue lines). (C-F) Structural mapping of conformational mobility profiles along first three slow modes for the RBD complexes with XGI-171, XGI-183, XGI-188 and XGI-203 antibodies. (G-J) Structural maps of conformational mobility profiles showing only RBD in surface colored according to the rigidity-to-flexibility scale : from blue (highly rigid) to red (highly flexible).

SCORE-B antibodies XGI-198 and XGI-203 target the RBM apex, overlapping the ACE2 interface. Their RMSF profiles show a dramatic rigidification of the RBM apex (residues 498–508) (Figure 3A). In XGI-198, residues T500, Y501, G502, V503, G504, H505, Q506, and Y508 exhibit RMSF values consistently below 0.7 Å, with Q506 as low as 0.26 Å. XGI-203 shows similarly low fluctuations in this region (T500, Y501, G502, V503) (Figure 3A). This rigid clamping effectively freezes the ACE2-binding surface, sterically blocking receptor engagement. In contrast, the RBM loop (470–490) remains flexible, with peaks reaching up to 3.5 Å in XGI-198 and 9 Å in XGI-203 (Figure 3A). This dynamic signature—a flexible periphery with a rigid core—explains why SCORE-B antibodies can tolerate peripheral mutations while maintaining high affinity for the conserved apex, and why XGI-198 achieves exceptional neutralization potency.

SCORE-C XGI-171 binds a highly conserved cryptic epitope on the inner face of the RBD. Its RMSF profile shows a remarkable dichotomy. The core epitope residues—Y369, F377, C379, Y380, T385—are strongly stabilized, with RMSF values below 0.8 Å, indicating that the antibody anchors firmly to this structurally invariant region (Figure 3A,B). In contrast, the RBM loop (470–490) exhibits dramatically elevated flexibility. Residues N481 and G482, which are part of the flexible β5-β6 loop, show RMSF values exceeding 4 Å (Figure 3A,B). This allosteric loosening is a direct consequence of antibody binding at the distal inner face; the interface does not directly contact the RBM, yet the binding induces a redistribution of rigidity that loosens the receptor-binding surface. The flexible RBM loop cannot engage ACE2 efficiently, explaining the weak neutralization of XGI-171 despite its ultra-broad binding. The RMSF profiles of CR3022 and EY6A, two well-characterized SCORE-C antibodies, mirror that of XGI-171 (Figure 3B). Both induce extreme flexibility in the RBM loop (470–490), with peaks exceeding 9 Å in CR3022 and 7 Å in EY6A. The core epitope residues remain rigid (RMSF < 1 Å). This consistent pattern across three structurally distinct antibodies targeting the same cryptic site confirms that SCORE-C antibodies function by allosterically loosening the RBM loop, impairing ACE2 binding without directly competing with the receptor.

Structural maps of conformational mobility illustrate this observation, showing contrasting rigidity of the binding epitope and elevated mobility of the RBM loop (Figure 3C,G). In fact, structural map of XGI-183 is very similar, where RBM loop (470–490) shows considerable flexibility (Figure 3D,H) that is higher than in SCORE-B antibodies. Structural maps for SCORE-B antibodies XGI-198 and XGI-203 illustrate striking rigidification of the RBD core and RBM apex (residues 498–508) (Figure 3E,F). The RMSF analysis across the three SCORE groups reveals a continuous spectrum of RBD flexibility modulation. SCORE-A antibodies stabilize the lateral epitope and moderately restrict the RBM loop, achieving a balance between breadth and potency. SCORE-B antibodies rigidly clamp the RBM apex, directly blocking ACE2. Their potent neutralization stems from locking the receptor-binding surface while tolerating peripheral loop mutations. SCORE-C antibodies induce maximal flexibility in the RBM loop, relying on allosteric destabilization to impair ACE2 binding. Their broad binding arises from rigid anchoring to a conserved core, but their weak neutralization reflects the indirect mechanism. Together, these data demonstrate that the conformational dynamics of the RBD–antibody interface are an integral part of the biophysical grammar that determines antibody function.

### 2.3. Mutational Profiling of Antibody-RBD Binding Interactions Interfaces Reveals Molecular Determinants of Immune Sensitivity

To provide a systematic comparison, we constructed mutational heatmaps for the RBD interface residues of the S complexes with XGI antibodies. Mutational scanning studies of RBD–antibody complexes provides insight into the epitope sensitivity and escape vulnerability of antibodies. For SCORE-A XG-183 antibody, mutational scanning revealed multiple strong hotspots T345, Y351, R355, K356, R357, F464, E465, R466, D467, I468, S469, T470, E471 (Figure 4A). Our results pointed to K356 as one of the major hotspots and experiments showed that XGI-183 had a diminished binding affinity towards BA.2.86, JN.1, KP.2 and KP.3 variants due to the K356T escape mutation (ΔΔG = 1.7 kcal/mol) which disrupts the salt bridge between K356 and D57 of XGi-183 (Figure 4). Despite targeting a similar epitope, S309 binding produced fewer hotspots with L335. P337, T345 and L441 are critical for S309 binding, with substitutions at these positions significantly impairing affinity (Supporting Information, Figure S1A). Another SCORE-A antibody SA58 revealed a number of similar hotspots including T345, L441, K444, V445 ((Supporting Information, Figure S1B). Despite simplified knowledge-based energetics of mutational scanning, these results are remarkably consistent with the experiments^44,53^ showing that full escape and strongest resistance are caused by mutations E340K, K444E while moderate resistance is caused by T345N and R346. Notably, weaker dependency on some vulnerable to mutations epitope residues may potentially be one of the reasons why S309 exhibits significantly stronger neutralizing activity [54] The analysis revealed that while these antibodies target a highly conserved core epitope, their neutralizing activity is constrained by specific mutation-induced disruptions.

**Figure 4.**
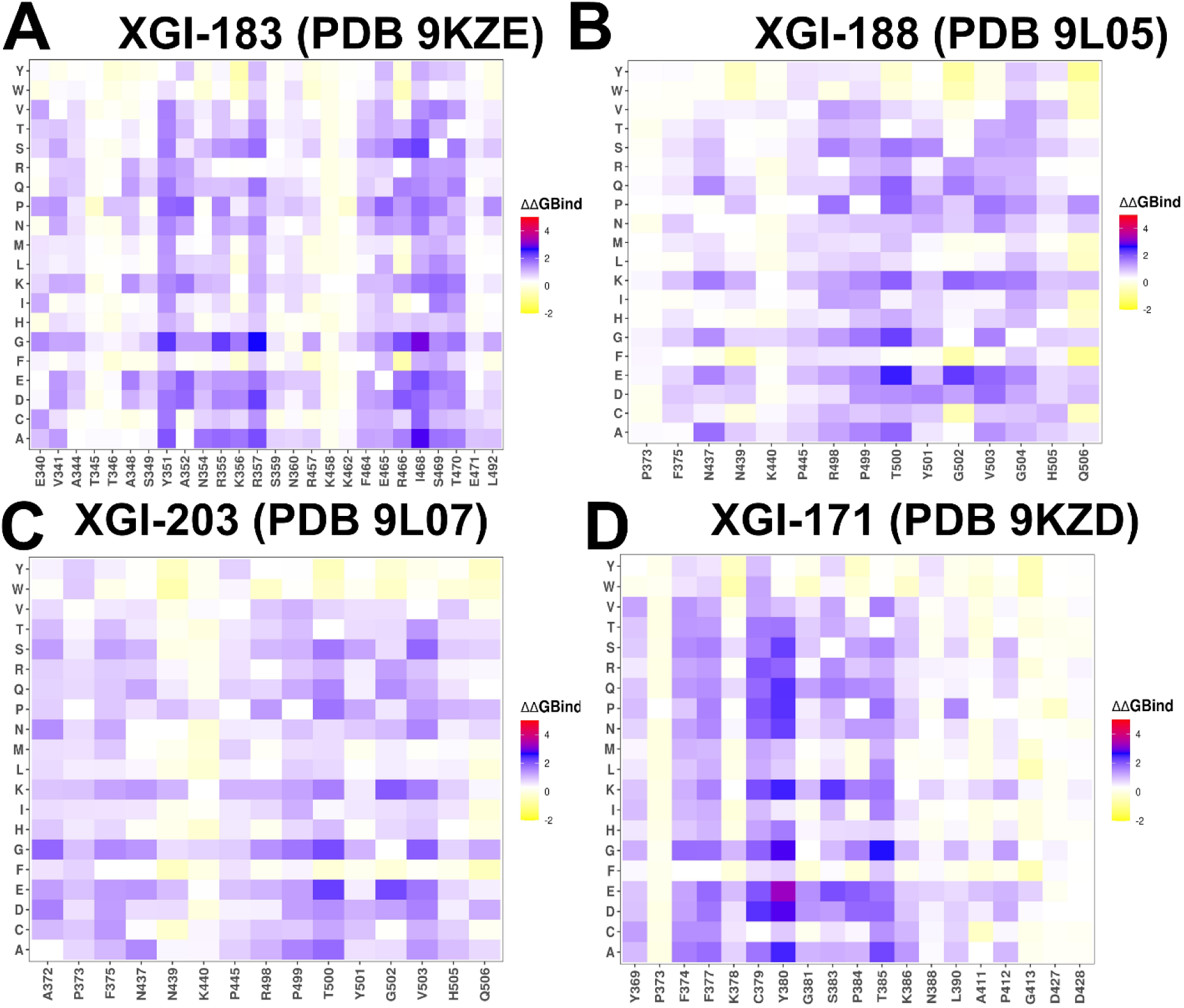
Mutational scanning of binding for the RBD complexes with XGI antibodies. The mutational scanning heatmaps for the binding epitope residues in the S-RBD complexes with XGI-183 with EG.5.1 RBD (A), XGI-188 with EG.5.1 RBD (B), and XGI-203 with EG.5.1 RBD (C) and XGI-171 with EG.5.1 RBD (D). The heatmaps show the computed binding free energy changes for 20 single mutations on the sites of variants. The squares on the heatmap are colored using a 4-colored scale blue-white-yellow-red, with blue indicating the largest unfavorable effect on binding and stability, while yellow-red points to mutations that have favorable effect and improve binding.

For SCORE-B XGI-198 primary hotspots are residues 498-500 and 503-506 (Figure 4B). Y505 and Q506 are highly conserved and are key ACE2 contacts; they form extensive hydrogen bonds and hydrophobic interactions with the antibody. The strong engagement of Y505 Q506 is unique to XGI-198 compared to XGI-203 (Figure 4C). Because XGI-198 potency relies heavily on the structural cross-linking mediated by Q506, a mutation at Q506 could compromise neutralization without fully abolishing binding [54]. However, the conservation of Q506 makes such escape rare. The virus can partially evade XGI-198 through mutations at peripheral residues (V503E, Y508H) without losing the core anchor.^54^ XGI-203 reliance on peripheral, less-constrained residues makes it the least potent and most escape-prone of the three. This hierarchy demonstrates that neutralization potency correlates with the number and functional importance of primary hotspots that directly overlap the ACE2 footprint. Mutational heatmap of SCORE-B antibody SA55 showed a greater number of strong hotspots including F374, S375, T376, D405, V407, R408, N437, G502, G504, and Y508 (Supporting Information, Figure S2). SA55 neutralizes more potently than XGI-198 and XGI-203 because its interface is anchored to a broader set of evolutionarily constrained ACE2-proximal residues, providing a direct and robust steric blockade. Among primary escape mutations identified via DMS studies are mutations V503A or G504A that can reduce SA55 binding [44,53]. For SCORE-C XGI-171 antibody the binding interface is dominated by a tightly packed hydrophobic cluster of binding hotspots formed by residues 374–385 of the RBD, with the strongest contributions from F377, C379, Y380, and T385 (Figure 4).

Structural mapping of binding hotspots across the three SCORE groups (Figure 5) provides a visual synthesis of the mutational scanning results. The spatial arrangement of hotspots—whether they cluster tightly around the ACE2 interface (SCORE-B), spread across lateral domains (SCORE-A), or bury deep within a cryptic pocket (SCORE-C)— provides structural rationale of the antibodies neutralization breadth. For SCORE-A antibodies, XGI-183, S309, and SA58 share a common hotspot core (K356, R357, T345, L441) but with distinct distributions: XGI-183 displays an extended network of hotspots that includes I468 and T470 on the β4-β5 hairpin, while S309 and SA58 show more localized hotspots concentrated around the α2-helix and the N343 glycan site (Figure 5A–C). In SCORE-B antibodies, XGI-198, XGI-203, and SA55 all anchor to the RBM apex, but their hotspot fingerprints differ markedly. XGI-198 uniquely engages Q506 and Y505 as dominant hotspots, creating a cross-linked clamp, whereas XGI-203 relies more heavily on T500 and V503, and SA55 broadens the hotspot repertoire to include F374, S375, T376, D405, V407, and R408 (Figure 5D–F). For SCORE-C antibodies, XGI-171, CR3022, and EY6A converge on a nearly identical hydrophobic core (F377, C379, Y380, T385), yet XGI-171 shows additional contributions from Y369 and K386, while CR3022 and EY6A extend hotspots to residues 383-386 and 390-392 (Figure 5G–I). This comparative mapping underscores a key principle: primary hotspots that are evolutionarily constrained and essential for antibody binding are conserved within each SCORE group, whereas secondary hotspots that are mutationally permissive vary among antibodies and can define their individual escape vulnerabilities.

**Figure 5.**
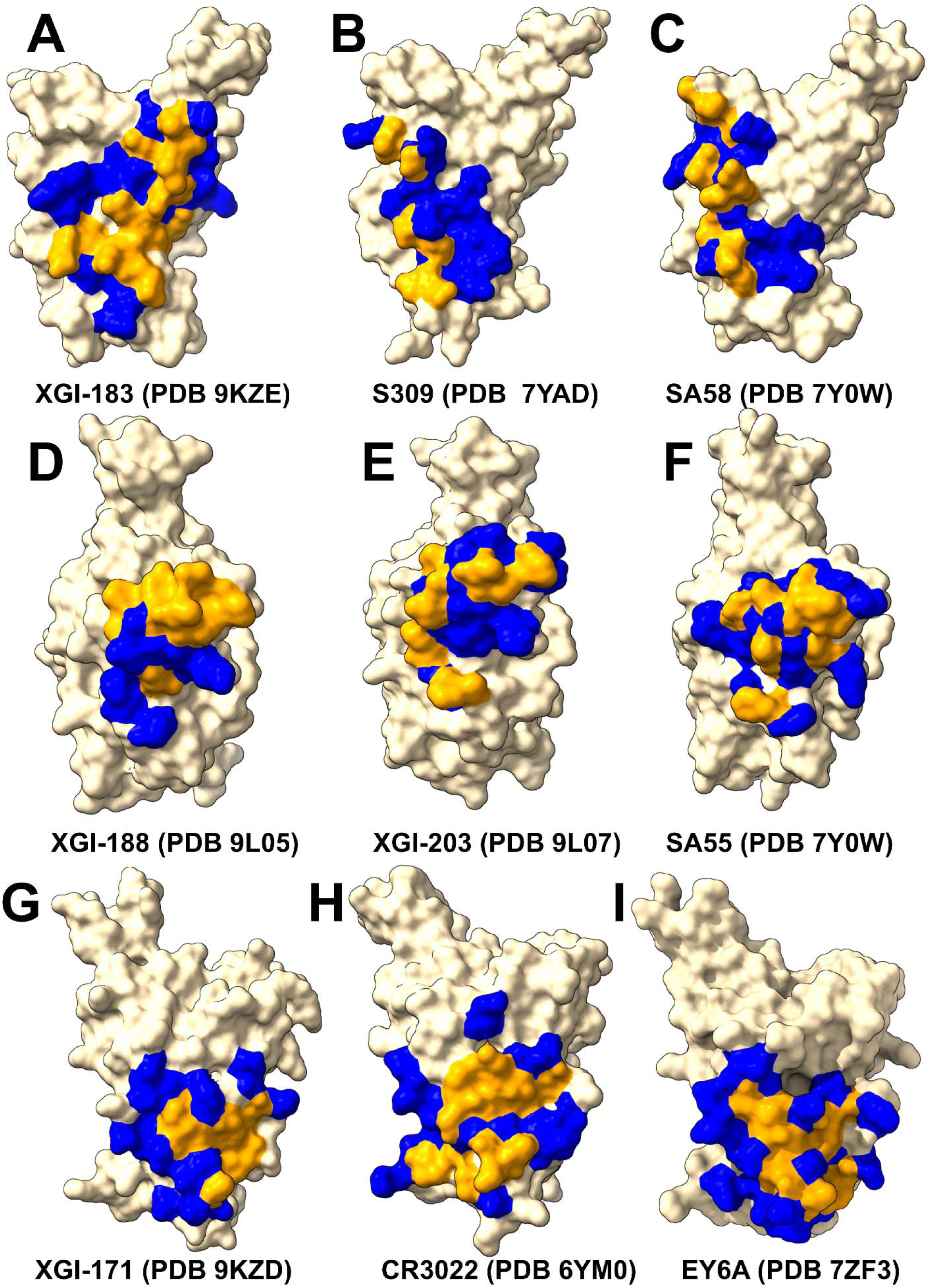
Structural Mapping of RBD Binding Hotspots for SCORE-A,B,C Antibodies Binding to the RBD. (A-C) The binding epitopes and binding hotspots are shown for three representative SCORE-A antibodies - XGI-183 (PDB ID: 9KZE), S309 (PDB ID: 7YAD), SA58 (PDB ID: 7Y0W). (D-F) The binding epitopes for three representative SCORE-B antibodies - XGI-188 (PDB ID: 9L05), XGI-203 (PDB ID: 9L07), SA55 (PDB ID:7Y0W). (G-H) The binding epitopes for three representative SCORE-C antibodies - XGI-171 (PDB ID:9KZD), CR3022 (PDB ID: 6YM0), EY6A (PDB ID: 7ZF3). The RBD is depicted in wheat-colored surface, with the heavy chain (HC) of the antibody in orange and the light chain (LC) in red. The binding epitope residues are shown in blue surface, the binding hotspots are in orange surface.

Due to the high level of sequence conservation of XGI-171-binding epitope across various sarbecoviruses, XGI-171 showed broad RBD binding activity for all the tested RBD or S proteins of SARS-CoV-2 variants [54] Mutational profiling of CR3022 binding identifies a set of key interface residues that are essential for antibody recognition. F377, C379, Y380, T385 emerged as dominant hotspots (Supporting Information, Figure S3A). Additional contributions come from K378, G381, V382, S383, P384, L390, and F392, which also play supportive roles in maintaining epitope integrity.

Our results are consistent with the DMS data revealing that residues 383–386, 390, and 392 represent major escape hotspots for CR3022. These residues lie at the edge of the epitope, suggesting that immune pressure drives mutations here to evade antibody recognition without compromising viral function. Mutational profiling revealed that EY6A engages similar dominant residues — including F377, C379, Y380, T385, and S383 — confirming that both antibodies recognize a shared structural motif (Supporting Information, Figure S3B). Crucially, none of the epitope residues overlap with the ACE2-binding motif. The mutational scanning data alone cannot explain weak neutralization by a loss of direct steric competition. Instead, the conformational dynamics data reveal that binding of SCORE-C antibodies induces allosteric loosening of the RBM loop (470–490), which becomes highly flexible (RMSF > 4 Å). This allosteric effect impairs ACE2 engagement indirectly, but it is inherently less efficient than direct receptor blockade. Because the neutralization mechanism relies on perturbing conformational equilibria rather than on a single high-affinity steric clash, even strong binding to the conserved core results in only weak neutralization. The mutational scanning data thus confirm the experiments [54] that the conserved epitope is highly resilient to escape, yet the allosteric mode of action explains the paradox of broad binding without potent neutralization.

In summary, mutational scanning across the three SCORE groups reveals a clear hierarchical organization of epitope sensitivity. For each antibody, a small set of primary hotspots—residues whose substitution consistently impairs binding—defines the energetic core of the interface. These primary hotspots are typically highly conserved and often overlap with ACE2-contact residues. Surrounding them are secondary hotspots, where mutations produce more moderate effects and which correspond to the sites of known immune-driven escape mutations in circulating variants (e.g., K356T for SCORE-A, V503E for SCORE-B, S383L for SCORE-C) [54]. The number and functional importance of primary hotspots directly correlate with neutralization potency: XGI-198 and SA55, which engage broader primary hotspot networks, show higher barriers to escape than XGI-203, which relies more heavily on secondary hotspots. For SCORE-C antibodies, the absence of ACE2-overlapping hotspots explains their weak neutralization despite a highly conserved epitope. Collectively, these mutational profiles provide a quantitative, residue-level map of antibody vulnerability and establish a predictive basis for anticipating which epitope positions are most likely to mutate under immune pressure.

### 2.4. Energetic Architecture of XGI Antibody–RBD Interfaces: Common Principles Across Three Conserved Epitopes

MM-GBSA analysis of the four XGI antibody–RBD complexes (Figure 6) together with the comparative analysis on related antibodies (S309, CR3022, EY6A, SA55), reveals a unifying energetic logic that underpins their broad neutralization and escape profiles. Across all three SCORE groups, the binding interfaces are organized around a conserved hydrophobic core that provides the primary energetic anchor, surrounded by peripheral residues whose energetic contributions are more modest and which are the principal sites of immune-driven mutations (Figure 6). However, the precise distribution of van der Waals (VDW) and electrostatic (ELE) forces determines the balance between potency and escape resistance. SCORE-A antibodies bind the lateral RBD through two distinct energetic clusters. The first is a VDW-dominated patch centered on residues I468 and T470. The second is an electrostatically reinforced cluster around K356 and R357 (Figure 6A-C). R357 emerges as the dominant hotspot, with strong electrostatic attraction (-57.97 kcal/mol) partially compensated by desolvation. Similarly, K356 makes strong contribution to the binding energy primarily due to electrostatic interactions (-64.92 kcal/mol) (Figure 6C). At the same time, I468 provides substantial hydrophobic stabilization due to burial in a hydrophobic pocket of the antibody paratope. Additionally, N360 and T345 contribute through polar interactions, consistent with antibody-antigen interfaces enriched in H-bond donors/acceptors (Figure 6A-C).

**Figure 6.**
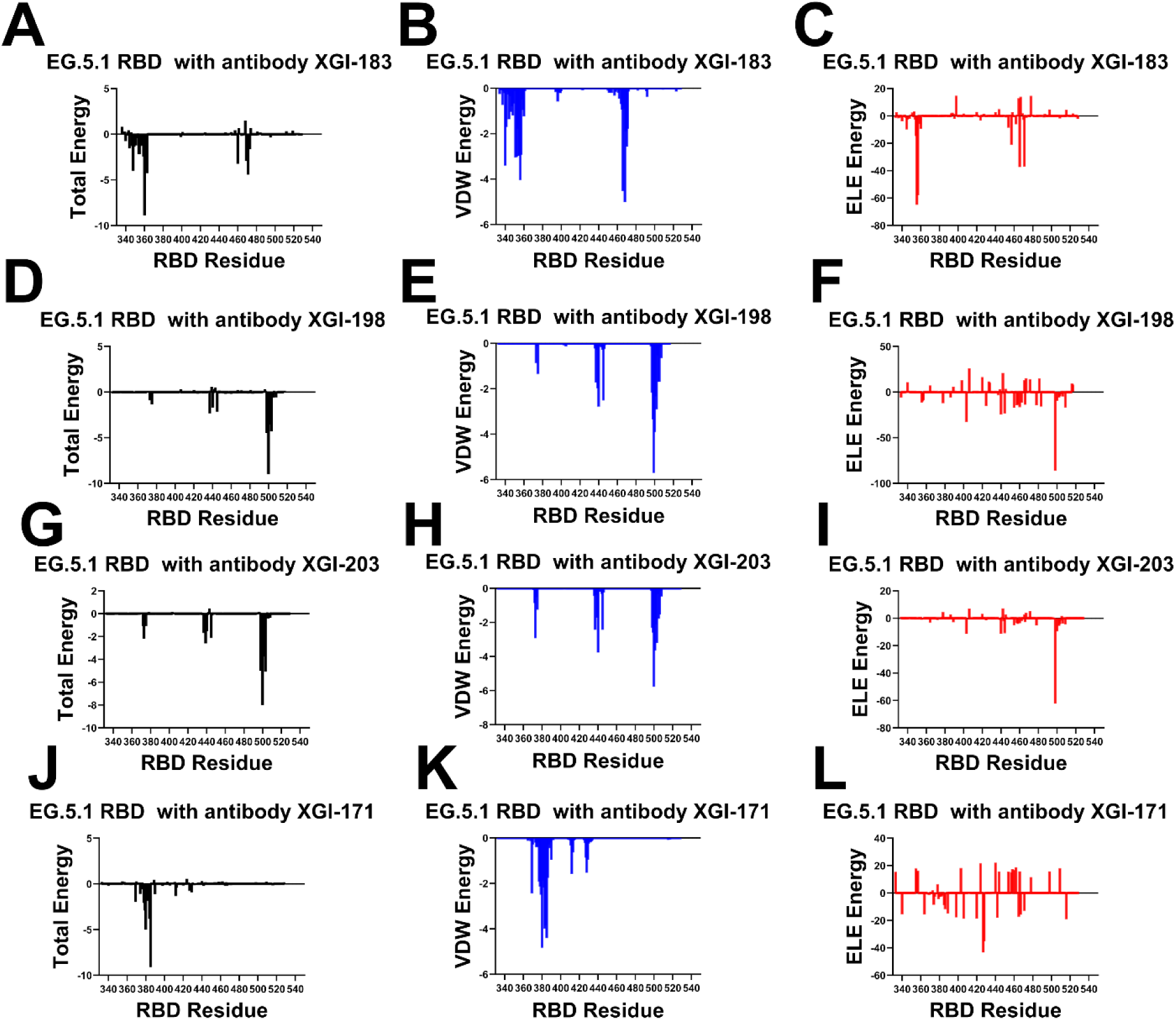
The residue-based decomposition of the binding MM-GBSA energies, van der Waals contributions and electrostatic interactions for the S-RBD complexes with XGI-183 (A-C), XGI-188 (D-F), XGI-203 (G-I) and XGI-171 (J-L). The binding free energy with MM-GBSA was computed by averaging the results of computations over 10,000 samples from the equilibrium ensembles. The standard error of the mean (SEM) for binding free energy estimates was calculated from the distribution of values obtained across the 10,000 snapshots sampled for each system. The statistical errors was estimated on the basis of the deviation between block average and are within 0.11-0.18 kcal/mol.

MM-GBSA analysis of S309 antibody complex shows a similar pattern, with K356 again appearing as a major electrostatic contributor, along with N343 and L441 (Supporting Information, Figure S4). The combined VDW and ELE contributions at K356 and R346 explain why mutations at these positions (K356T, R346K) have repeatedly emerged to escape S309. The residue decomposition of the total binding energy for S309 (Supporting Information, Figure S4) revealed strong and consistent binding hotspots for T345(ΔG = -4.72 kcal/mol), K356 (ΔG = - 3.29 kcal/mol), L441 (ΔG = -2.83 kcal/mol) N343 (ΔG = -3.03 kcal/mol), A344 (ΔG = -1.72 kcal/mol), and R346 positions (ΔG = -1.69 kcal/mol) (Supporting Information, Figure S4). This is consistent with mutational scanning computations that identified these residues as critical for binding S309 antibody. The MM-GBSA residue decomposition analysis further underscores the critical role of L441 and K444 in stabilizing the S309-RBD complex. The energy decomposition showed that the strongest van der Waals interactions are provided by T345, P337, A344, L335 and L441 RBD residues where T345 and P337 are key hotspots that are driven by favorable hydrophobic contacts (Supporting Information, Figure S4B).

SA58 relies on a mixed electrostatic/van der Waals architecture similar to XGI-183, with K444 contributing favorable electrostatic interactions (analogous to K356 in XGI-183) and L441 providing hydrophobic stabilization (Supporting Information, Figure S5). However, its weaker dependence on K356 (which is not a major hotspot for SA58) and its engagement of K444 and R509 suggest a shifted energetic center that may confer different escape sensitivity. Thus, SCORE-A antibodies achieve moderate potency through a dual-anchor strategy, but their dependence on a few electrostatically sensitive residues K356 makes them vulnerable to escape via mutations that disrupt those interactions.

SCORE-B antibodies target the RBM apex, and their binding is overwhelmingly driven by VDW interactions with residues T500, V503, R498, G502, and N437. In XGI-188, the top hotspots are T500 (ΔG = -8.98 kcal/mol), R498 (ΔG = -4.47 kcal/mol), V503 (ΔG = -4.29 kcal/mol), and G502 (ΔG = -3.57 kcal/mol) (Figure 6D-F). For XGI-203, the key binding hotspots are essentially the same, T500 (ΔG = -8.0 kcal/mol), V503 (ΔG = -5.09 kcal/mol) and R498 (ΔG = -5.01 kcal/mol) (Figure 6G-I). Electrostatic contributions are modest and largely compensated, indicating that the interface is shaped primarily by shape complementarity and hydrophobic packing. The conserved core residues (T500, V503, R498, G502) are under strong purifying selection because they are essential for ACE2 binding. This explains the high barrier to escape for SCORE-B antibodies: mutations at these positions would cripple viral fitness. Peripheral residues like V503 and Y508 have been observed to mutate (e.g., V503E, Y508H) without abolishing binding, consistent with their secondary-hotspot status (ΔΔG 1.9–2.2 kcal/mol). MM-GBSA of SA55 binding showed similar hotspots primarily Y501 (ΔG = - 5.56 kcal/mol) and T500 (ΔG = -4.86 kcal/mol) with major contribution of van der Waals interqc4tions (Supporting Information, Figure S6). Thus, SCORE-B antibodies combine a VDW-dominated, evolutionarily constrained core with a structurally critical peripheral interaction that unlocks superior potency.

For SCORE-C antibodies (XGI-171, CR3022, EY6A), the binding interface is dominated by a tightly packed hydrophobic cluster formed by residues 369–385 of the RBD, with the strongest contributions coming from Y369, F377, C379, Y380, and T385. In XGI-171, T385 alone contributes nearly −9 kcal/mol to the total binding energy—more than any other single residue in the complex—and this contribution is roughly equally split between van der Waals and electrostatic interactions, reflecting its dual role as both a hydrophobic anchor and a hydrogen-bond partner. The peripheral residues S383, P384, and K386 contribute less than −2 kcal/mol each, which is consistent with their classification as secondary hotspots in mutational scanning (ΔΔG 1.5–2.2 kcal/mol). This energetic hierarchy explains the ultra-broad binding of XGI-171: the core is so strongly anchored that even when the periphery mutates (e.g., S383L, T385I in Omicron), the interface remains intact.

CR3022 binds the RBD through a hydrophobic core centered on residues F377, C379, Y380, and T385. The total binding energy decomposition shows that these residues contribute the most favorable van der Waals interactions, while electrostatic contributions are modest and largely offset by desolvation penalties (Supporting Information, Figure S7). Notably, residues S383, P384, K386, L390, and F392 provide secondary stabilization, consistent with their identification as escape hotspots in deep mutational scanning studies. Compared to XGI-171, CR3022 shows a broader distribution of energetic contributions across the 383-392 region, which may explain why mutations at S383L or T385I partially reduce binding without completely abolishing it. The absence of significant electrostatic anchoring makes CR3022 less sensitive to polar substitutions but also less potent overall. EY6A engages a similar hydrophobic core but extends its interaction to residues P412 and G413 on the β7 strand. The MM-GBSA decomposition for EY6A shows that van der Waals interactions again dominate, with F377, C379, Y380, and T385 as primary contributors. However, the additional contacts with P412 and G413 provide supplementary hydrophobic stabilization which may confer resilience against certain mutations in the 383-386 region. Interestingly, electrostatic contributions from K378, K386, D405, D427, and D428 are largely compensated by unfavorable solvation terms, indicating that EY6A relies almost exclusively on shape complementarity and hydrophobic packing (Supporting Information, Figure S8). This energetic profile explains why EY6A retains binding to many variants that escape CR3022, but also why its neutralization potency remains weak. Taken together, the MM-GBSA profiles of CR3022 and EY6A confirm the SCORE-C paradigm: a deeply buried, minimally frustrated hydrophobic core provides ultra-broad binding, while peripheral residues (383-386, 390-392, and, in EY6A, 412-413) contribute modestly and serve as secondary escape hotspots. The convergence of these energetic signatures across three structurally distinct antibodies (XGI-171, CR3022, EY6A) validates the robustness of the SCORE-C epitope as an evolutionarily constrained anchor, while the subtle differences in peripheral contacts explain the graded escape resistance observed experimentally.

To summarize, MM-GBSA profiles across the three SCORE groups reveal a consistent organizing principle: broad neutralization is achieved by anchoring binding to evolutionarily constrained cores, while potency is modulated by strategically placed electrostatic or structural peripheries. SCORE-A antibodies rely on a dual hotspot architecture; their vulnerability lies in electrostatically sensitive residues like K356, which have become hotspots for immune escape (K356T). SCORE-B antibodies are dominated by van der Waals interactions with the RBM apex. SCORE-C antibodies anchor to a deeply buried hydrophobic core, ensuring ultra-broad binding. The hotspots identified here align precisely with the mutational scanning data on primary hotspots and this convergence validates the use of MM-GBSA as a predictive tool for mapping antibody escape pathways. Our results suggest that broad neutralization is achieved by anchoring to evolutionarily constrained cores, while potency is modulated by strategically placed electrostatic or structural peripheries. MM-GBSA hotspots align precisely with mutational scanning data, validating this approach for predicting antibody escape pathways.

### 2.5 Frustration Landscape Analysis of Antibody–RBD Interfaces Reveals Energetic Signatures of Binding and Resistance to Immune Escape

To dissect the energetic determinants of antibody binding and immune escape, we computed both conformational frustration (sensitivity to structural perturbations) and mutational frustration (sensitivity to amino acid substitutions) for four XGI antibodies in complex with the SARS-CoV-2 EG.5.1 spike RBD. This analysis maps residues as minimally frustrated (evolutionarily optimized and rigid), neutrally frustrated (plastic and mutation-tolerant), or highly frustrated (strained and conformationally labile), providing a high-resolution view of the energetic architecture at antibody–antigen interfaces. The frustration density profiles across RBD residues 330–530 reveal distinct architectural patterns that correlate with each antibody’s binding mode and epitope location (Figure 7). For the SCORE A antibody XGI-183, the conformational frustration landscape displays a heterogeneous pattern with prominent neutral frustration peaks interspersed with regions of minimal frustration (Figure 7A). Elevated neutral frustration density appears in two discrete regions—residues 350–360 and 460–480—corresponding to the lateral epitope characteristic of SCORE A antibodies. Sharp peaks of highly frustrated residues appear at specific positions, notably near residues 357 and 468, marking potential conformational switch points that may serve as initial binding sensors. The mutational frustration profile for XGI-183 reveals broader regions of minimal frustration at residues 340, 440, and 500, indicating evolutionarily constrained positions (Figure 7B). Critically, neutral frustration dominates precisely the central epitope regions (350–370 and 460–480), suggesting these positions tolerate substitutions—an observation consistent with known escape mutations at K356 and R357.

**Figure 7.**
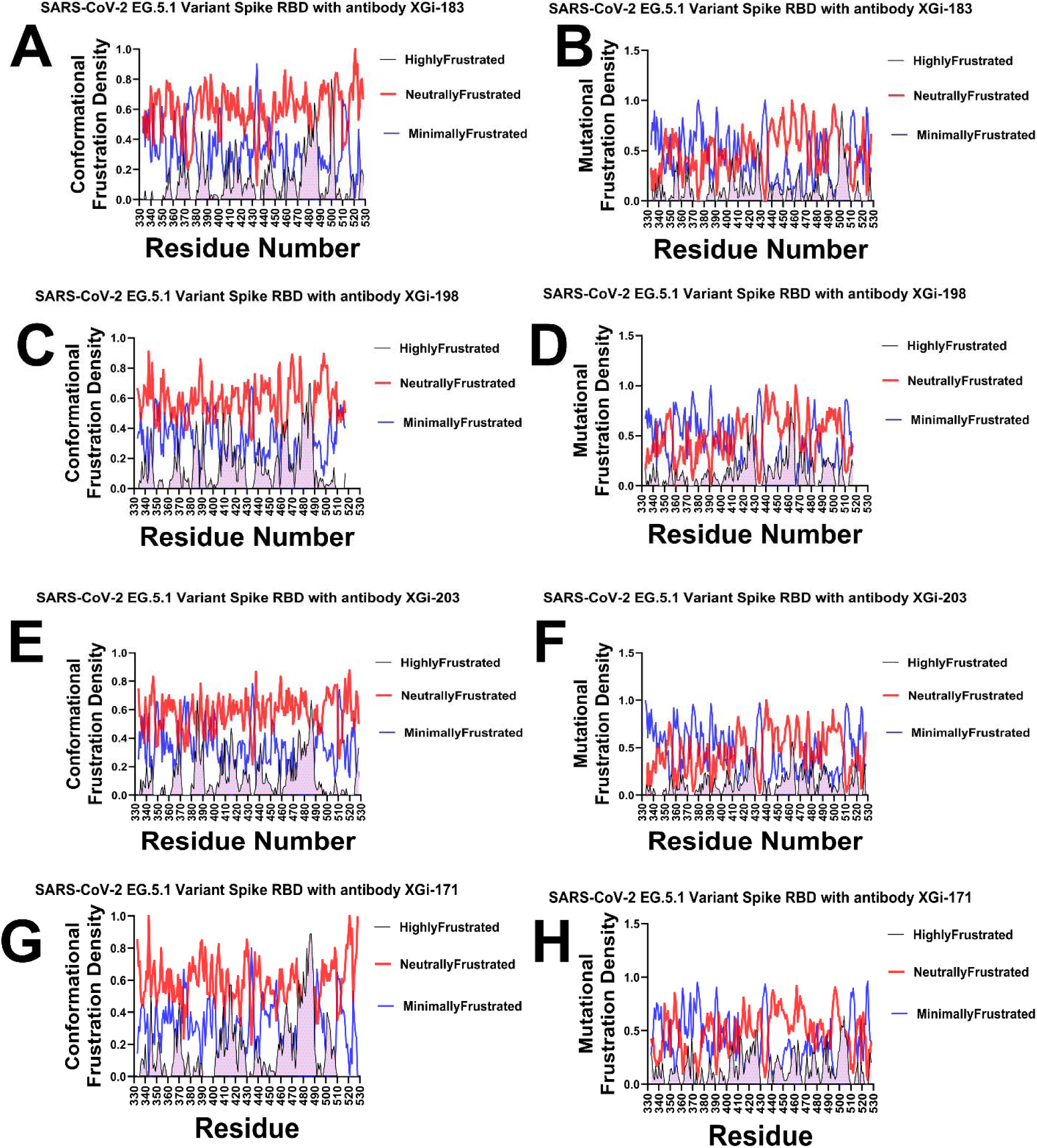
Frustration density profiles along the RBD sequence for XGI antibody complexes. (A–H) Local frustration density (Z-score) mapped onto RBD residues 330–530 for four XGI antibodies in complex with the SARS-CoV-2 EG.5.1 spike RBD. Conformational frustration (left column, panels A, C, E, G) measures sensitivity to structural perturbations; mutational frustration (right column, panels B, D, F, H) measures sensitivity to amino acid substitutions. Residues are colored according to the dominant frustration class: red = neutrally frustrated (plastic, mutation-tolerant), blue = minimally frustrated (evolutionarily optimized, rigid), green = highly frustrated (strained, conformationally labile). (A, B) SCORE-A antibody XGI-183: neutral frustration dominates the lateral epitope (350-360, 460-480), consistent with escape mutations at K356 and R357; minimal frustration appears at positions 340, 440, and 500. (C, D) SCORE-B antibody XGI-198: similar to XGI-183 but with increased minimal frustration around residues 440-460 and residue 500, indicating stronger evolutionary constraint. (E, F) SCORE-B antibody XGI-203: conformational neutral frustration spans the RBM apex (470-500), while mutational frustration shows sharp minimal-frustration peaks at the ACE2-binding ridge (498-502), explaining its high escape barrier. (G, H) SCORE-C antibody XGI-171: a prominent minimal-frustration peak centers on the cryptic epitope (475-490) in both profiles, with highly frustrated residues at the periphery, marking the signature of ultra-broad binding. These profiles demonstrate that the balance among frustration classes, rather than any single class, defines epitope architecture and escape vulnerability.

The SCORE A antibody XGI-198 shows subtle shifts in frustration density (Figure 7C, D). The neutral frustration peaks in the 340–360 region are slightly more pronounced than in XGI-183, while minimal frustration increases around residue 500. This shift suggests XGI-198 engages a subtly shifted epitope with altered energetic constraints. The mutational profile reinforces this interpretation, showing increased minimal frustration around residues 440–460 compared to XGI-183, indicating stronger evolutionary constraint at these positions that may reflect optimized contacts reducing escape potential (Figure 7D).

XGI-203 exhibits further gradual shifts in the frustration architecture (Figure 7E, F). Conformationally, it shows prominent neutral frustration across the receptor-binding motif (RBM) apex spanning residues 470–500. Rather than appearing as sharp peaks, this region displays sustained neutral frustration density, suggesting a broad, plastic interface characteristic of RBM apex targeting (Figure 7E). The mutational profile reveals a critical finding: strong minimal frustration peaks at residues 498–502, corresponding to the ACE2-binding ridge (Figure 7F). This indicates that SCORE B antibodies target evolutionarily constrained positions essential for viral fitness, explaining their high barrier to escape. SCORE C antibody XGI-171 displays the most distinctive frustration profile of all four antibodies (Figure 7G, H). The conformational frustration landscape features a prominent minimal frustration peak centered at residues 475–490, corresponding to the cryptic, conserved epitope characteristic of SCORE C antibodies (Figure 7G).

Highly frustrated peaks appear at the periphery near residues 350 and 520, marking regulatory switch points. The mutational profile shows the strongest minimal frustration signal of all four antibodies, particularly concentrated around residues 370–385 and 475–490 (Figure 7H). This indicates that XGI-171 targets the most evolutionarily constrained regions of the RBD, consistent with its ultra-broad binding across all variants.

Together, the frustration density profiles establish a clear gradient in energetic architecture across the three SCORE classes. The SCORE-A antibody XGI-183 displays a mixed landscape: neutral frustration marks the central epitope regions (350-370 and 460-480), indicating mutational permissiveness and explaining its vulnerability to escape mutations such as K356T and R357T. Minimal frustration appears in flanking zones (340, 440, 500), reflecting evolutionary constraint at positions not directly targeted by the antibody. The two SCORE-B antibodies, XGI-198 and XGI-203, show progressively more constrained architectures. XGI-198 shifts toward increased minimal frustration near residues 440-460 and residue 500, suggesting a modest gain in escape resistance compared to XGI-183. XGI-203 exhibits a distinctive decoupling: conformational neutral frustration spans the broad, plastic RBM apex (470-500), while mutational frustration reveals sharp minimal-frustration peaks at the ACE2-binding ridge (498-502). Finally, the SCORE-C antibody XGI-171 exhibits the most extreme profile, with a prominent minimal-frustration peak centered on the cryptic epitope (475-490) in both conformational and mutational maps, alongside neutral frustration elsewhere. Together, these profiles demonstrate that the balance between neutral, minimal, and highly frustrated regions—not simply the presence of any single class—dictates an antibody’s functional breadth and resilience.

#### Comparative Interface-Specific Frustration Distributions: Conformational Plasticity Versus Mutational Constraint

To quantify the energetic organization at the binding interface and to compare how conformational flexibility and mutational tolerance are distributed across the binding epitope, we computed frustration density distributions specifically for RBD residues within 4 Å of antibody heavy atoms. These distributions reveal a unifying architectural principle across all four XGI antibodies: neutral frustration dominates the binding interface, while highly and minimally frustrated residues play specialized, complementary roles. However, critical differences emerge when comparing conformational versus mutational frustration distributions, illuminating the distinct biophysical mechanisms that govern binding affinity versus resistance to immune escape (Figure 8). For XGI-183, the conformational frustration distribution shows a characteristic pattern where neutral frustration dominates at high relative frustration densities in the 0.5–0.8 range (Figure 8A, left). Highly frustrated residues contribute primarily at low relative densities (0.0–0.2), indicating they represent only a small fraction of the interface and are likely localized to specific switch points rather than distributed across the epitope. Minimally frustrated residues show modest contributions across mid-range densities (0.2–0.4), suggesting a limited number of evolutionarily optimized contacts. The mutational distribution (Figure 8A, right) reinforces this architecture but with notable shifts: neutral frustration dominates across a slightly lower density range (0.4–0.7), while highly frustrated residues peak even more sharply at very low densities (0.0–0.1). This discrepancy between the conformational and mutational profiles is instructive. The conformational distribution indicates that the interface possesses moderate structural flexibility, with neutral frustration providing plasticity for induced-fit adjustments. However, the mutational distribution reveals that this same interface is highly permissive to amino acid substitutions—the neutral frustration in the mutational landscape marks positions that can be altered without compromising the overall fold or binding competence (Figure 8B). This duality explains the observed escape at K356 and R357: these residues are conformationally neutral (allowing structural adaptation during binding) but mutationally neutral (allowing sequence changes that ablate antibody recognition without fitness cost).

**Figure 8.**
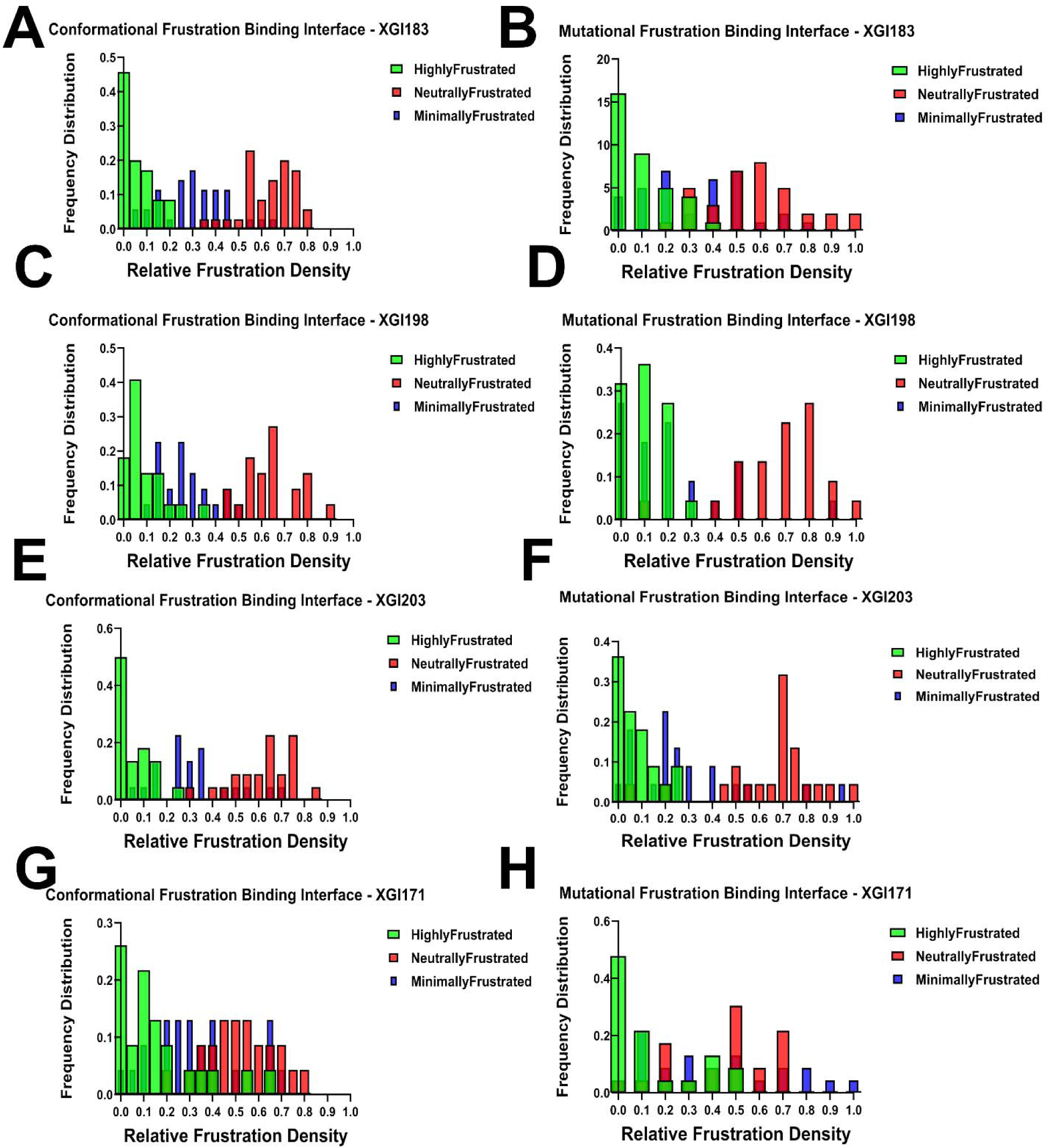
Interface-specific frustration distributions for XGI antibody–RBD binding interfaces. Relative frustration density distributions computed exclusively for RBD residues within 4 Å of antibody heavy atoms. For each antibody, the left panel of the pair shows conformational frustration and the right panel shows mutational frustration. Bars are colored as red = neutrally frustrated, blue = minimally frustrated, green = highly frustrated. (A, B) XGI-183 (SCORE-A): Neutral frustration dominates at high relative densities (0.5-0.8) in both distributions, while minimally frustrated residues contribute modestly at mid-range densities. (C, D) XGI-198 (SCORE-B): Neutral frustration remains dominant, but the mutational peak shifts to a higher density (∼0.75), and highly frustrated contributions increase slightly in the conformational distribution (0.0-0.2). This subtle decoupling reflects reduced mutational tolerance compared to XGI-183, conferring improved escape resistance while retaining a neutral-frustration scaffold. (E, F) XGI-203 (SCORE-B): The conformational distribution shows the highest highly frustrated contribution of the four antibodies (green peak at 0.0-0.1), yet neutral frustration still dominates at higher densities (0.5-0.7). The mutational distribution diverges markedly: minimally frustrated residues contribute substantially at 0.2-0.3 density, and neutral frustration peaks at 0.6-0.8. (G, H) XGI-171 (SCORE-C). Minimally frustrated residues make substantial contributions at high relative densities (0.7-0.9) in both conformational and mutational distributions—a pattern unique to this antibody. Highly frustrated residues dominate at low densities (0.0-0.2).

XGI-198 displays a similar neutral frustration-dominated pattern in its conformational distribution, with red bars prominent at 0.5–0.7 densities (Figure 8C). However, highly frustrated contributions are slightly elevated compared to XGI-183, particularly in the 0.0–0.2 range, suggesting XGI-198 introduces more conformational strain at the interface while maintaining the neutral frustration scaffold. The mutational distribution (Figure 8D) shows a prominent neutral frustration peak shifted to a higher density at approximately 0.75, with highly frustrated residues concentrated at low densities. Critically, the mutational neutral frustration peak appears at higher relative density than in XGI-183, indicating that the residues engaged by XGI-198 are less mutationally tolerant—they reside in a region of the energy landscape where sequence changes are more likely to disrupt favorable interactions. This pattern reveals a key mechanistic insight: XGI-198 achieves improved affinity not by eliminating neutral frustration but by shifting the balance toward residues that are mutationally less permissive. The antibody balances strain (introduced through contacts with highly frustrated residues) with plasticity (maintained through the neutral frustration scaffold) while simultaneously reducing the mutational accessibility of its epitope. This explains why XGI-198 shows greater escape resistance than XGI-183 despite belonging to the same SCORE A class.

XGI-203 exhibits the highest highly frustrated contribution of the four antibodies in its conformational distribution, with green peaks at 0.0–0.1 (Figure 8E), yet neutral frustration still dominates at higher densities (0.5–0.7). This indicates XGI-203 engages a substantial number of conformationally strained, sensitive positions—consistent with RBM apex targeting where structural precision is required for ACE2 mimicry. The mutational distribution (Figure 8F) reveals a distinctive pattern that diverges markedly from the conformational profile: while neutral frustration dominates at 0.6–0.8 densities, minimally frustrated residues contribute more substantially than in XGI-183 or XGI-198, particularly around 0.2–0.3 density. The divergence between the conformational and mutational distributions for XGI-203 is the most pronounced among the four antibodies.

Conformationally, the interface appears highly plastic, with sustained neutral frustration across the RBM apex suggesting broad adaptability. Mutationally, however, the interface reveals a constrained core where minimally frustrated residues—positions that are evolutionarily optimized and intolerant to change—make significant contributions. This apparent paradox resolves when considering the biological function of the RBM apex: residues 498–502 form the ACE2-binding ridge, a region under strong purifying selection because any mutation impairs receptor binding. XGI-203 targets precisely this region. The conformational neutral frustration reflects the structural plasticity required for the RBD to adopt the “up” conformation and engage ACE2, while the mutational minimal frustration reflects the evolutionary constraint that makes these positions invulnerable to escape mutations. Thus, XGI-203 achieves its high escape barrier by anchoring its interface to mutationally constrained residues, even though the surrounding structural scaffold remains conformationally plastic.

XGI-171 displays a qualitatively distinct architecture that sets it apart from the other three antibodies (Figure 8D). In the conformational distribution, while neutral frustration remains prominent at 0.4–0.6 densities, minimally frustrated residues make substantial contributions at high relative densities in the 0.7–0.9 range—a pattern not observed elsewhere. Highly frustrated residues dominate at low densities (0.0–0.2), as expected. The mutational distribution reveals the most striking feature: minimally frustrated residues contribute significantly at high relative densities above 0.7, alongside neutral frustration peaks at 0.5–0.6. XGI-171 is the only antibody in which minimally frustrated positions substantially populate the binding interface in both conformational and mutational distributions, and crucially, the two distributions align closely with each other. This convergence indicates that for XGI-171, conformational rigidity and mutational constraint coincide at the same set of residues. The cryptic epitope targeted by SCORE C antibodies is not only evolutionarily locked (mutational minimal frustration) but also structurally rigid (conformational minimal frustration), creating an interface that offers no easy escape routes. The virus cannot mutate these residues without losing fitness, nor can it adopt alternative conformations to evade binding.

Together, these distributions reveal that the relationship between conformational and mutational frustration—whether aligned (XGI-183), subtly decoupled (XGI-198), strongly decoupled (XGI-203), or convergently minimal (XGI-171)—encodes distinct mechanisms of binding, potency, and immune escape across the three SCORE classes. The comparative analysis across all four antibodies reveals a fundamental principle: the relationship between conformational and mutational frustration distributions encodes distinct binding mechanisms. Antibodies like XGI-183 show aligned distributions where neutral frustration dominates in both, yielding permissive interfaces that are plastic but escape-prone. XGI-198 shows a subtle decoupling, with mutational neutral frustration shifted to higher densities, indicating a moderate increase in escape resistance without fundamentally altering the interface architecture. XGI-203 displays pronounced decoupling, where conformational neutral frustration coexists with mutational minimal frustration, enabling targeting of functionally essential regions while maintaining structural adaptability. XGI-171 shows convergence, where minimal frustration dominates in both distributions, creating an interface that is both structurally rigid and evolutionarily locked—the signature of ultra-broad neutralization.

## 3. Discussion

The continued evolution of SARS-CoV-2 has rendered most clinically approved monoclonal antibodies ineffective, yet a small number of broadly neutralizing antibodies targeting three newly identified super-conserved RBD epitopes—SCORE-A, SCORE-B, and SCORE-C—retain remarkable activity against even the most recent JN.1-derived sublineages. The systematic characterization of XGI antibodies by Cao and colleagues [54] revealed three non-overlapping, super-conserved RBD epitopes—SCORE-A, SCORE-B, and SCORE-C—that remain accessible despite extensive antigenic drift. However, the structural and energetic principles that endow these epitopes with differential neutralization potency and escape resistance remained unclear. To investigate these questions, we employed a multi-pronged computational framework integrating conformational dynamics, mutational scanning, MM-GBSA binding energetics, and frustration profiling to dissect the molecular mechanisms by which XGI antibodies achieve broad neutralization and resistance to immune escape. Our results reveal a unifying biophysical architecture: all three SCORE epitopes are organized around a highly conserved, minimally frustrated core that provides stable anchoring, flanked by peripheral regions whose energetic and dynamic properties determine neutralization potency and escape vulnerability.

SCORE-A antibodies (exemplified by XGI-183) bind the lateral RBD surface, rigidifying the α2-helix and β4-β5 hairpin while leaving the RBM loop partially mobile. Their binding energy is distributed between a van der Waals-dominated patch (I468, T470) and an electrostatically reinforced cluster (K356, R357). The frustration profiles show that the central epitope regions (350-370, 460-480) are neutrally frustrated, explaining why mutations at K356 and R357 are tolerated by the virus and have emerged as escape hotspots. Taken together, the synergy of methods reveals a complete mechanism: XGI-183 achieves moderate potency by rigidifying a lateral anchor while relying on electrostatic hotspots that are energetically important but evolutionarily dispensable. The virus escapes by mutating these neutrally frustrated positions (K356T, R357T), which reduces antibody affinity without sacrificing viral fitness. Our results suggest that the permissive neutral-frustration scaffold, identified through frustration analysis may be the root cause of the observed escape vulnerability [54]. SCORE-B antibodies (XGI-198 and XGI-203) target the RBM apex, directly blocking ACE2. Their binding is overwhelmingly driven by van der Waals interactions with evolutionarily constrained residues (T500, V503, R498, G502). The dynamic signature is a rigid clamp at the ACE2-binding ridge combined with a flexible RBM loop. Frustration analysis reveals a striking decoupling: conformational neutral frustration across the broad RBM apex enables structural adaptability, while mutational minimal frustration at the ACE2-binding ridge makes these positions invulnerable to escape mutations. The frustration analysis reveals that Q506, while conserved, is not as deeply minimally frustrated as T500 or R498, explaining why even XGI-198 has some residual escape vulnerability (e.g., via V503E or Y508H). Thus, the intragroup gradient is directly attributable to the number and functional importance of primary hotspots that overlap the ACE2 footprint. Our results established a potential mechanism in which XGI-198 achieves superior potency and resistance because it engages a broader set of minimally frustrated, ACE2-proximal hotspots (including Q506 and Y505), whereas XGI-203 relies more on neutrally frustrated peripheral residues.

XGI-171 achieves ultra-broad binding because it anchors to a minimally frustrated core that the virus cannot mutate without losing fitness. However, its weak neutralization arises because the mechanism is allosteric—destabilizing the RBM loop indirectly—rather than directly blocking ACE2. The allosteric effect, while sufficient to impair receptor engagement, is inherently less efficient than a direct steric clash. Frustration analysis confirms that none of the epitope residues overlap with the ACE2-binding motif, explaining the absence of direct competition. Thus, the experimental observation of broad binding but weak neutralization is not a contradiction but a direct consequence of the allosteric, minimally frustrated architecture revealed by our multi-method approach.

Across all three SCORE classes, the integration of dynamics, mutational scanning, MM-GBSA, and frustration analysis reveals a consistent organizing principle: broad neutralization arises from anchoring to minimally frustrated cores, while neutral frustration defines the accessible escape pathways. Our findings directly explain the experimental observations of Cao and colleagues [54] why XGI-183 is escape-prone (neutral frustration at K356/R357), why XGI-198 is more potent than XGI-203 (broader primary hotspot network), and why XGI-171 binds everything but neutralizes weakly (allosteric mechanism from a minimally frustrated core). More broadly, this framework transforms our understanding of antibody durability. Rather than seeking maximally high affinity, we should design antibodies that distribute binding energy across minimally frustrated, evolutionarily constrained cores. Frustration profiling can prospectively rank epitopes and antibodies, enabling the selection of candidates that target the virus’s invulnerable centers. The pattern of “energetically suboptimal, adaptable frustration” is not a limitation of viral adaptability—it is the very feature that makes broad and durable immune responses possible. These insights may help to transform how we think about antibody-driven immune pressure. Rather than viewing escape as an unpredictable arms race, we can now anticipate the most likely escape mutations by mapping the neutral frustration landscape of the RBD. For antibody discovery, this suggests that the most durable antibodies will be those that engage minimally frustrated residues as their primary anchors, while tolerating—or even exploiting—the predictable mutational plasticity of neutrally frustrated peripheries. By deliberately targeting the invulnerable core, such antibodies create a high barrier to resistance; any escape that does occur is confined to peripheral residues that provide only partial relief, allowing the antibody to retain activity for longer. Moreover, this framework informs vaccine design: immunogens that focus the immune response on minimally frustrated, functionally indispensable regions of the RBD may elicit antibodies with a higher genetic barrier to escape. By understanding the frustration landscape of the RBD, we can design next-generation vaccines that steer the immune system toward the stable, evolutionarily constrained sites that the virus cannot afford to mutate.

## 3. Materials and Methods

### 3.1 Structure Preparation and Analysis

All structures were obtained from the Protein Data Bank [81]. Hydrogen atoms and missing residues were initially added and assigned according to the WHATIF program web interface [82,83]. The structures were further pre-processed through the Protein Preparation Wizard (Schrödinger, LLC, New York, NY) for assignment and adjustment of ionization states, formation of assignment of partial charges as well as additional check for possible missing atoms and side chains that were not assigned by the WHATIF program. The missing loops in the cryo-EM structures were reconstructed using template-based loop prediction approaches ModLoop [84] and ArchPRED [85] and further confirmed by FALC (Fragment Assembly and Loop Closure) program [86]. The side chain rotamers were refined and optimized by SCWRL4 tool [87]. The protein structures were then optimized using atomic-level energy minimization using 3Drefine method [88].

### 3.2 Coarse-Grained Simulations

We employed CABS-flex approach that efficiently combines a high-resolution coarse-grained model and efficient search protocol capable of accurately reproducing all-atom MD simulation trajectories and dynamic profiles of large biomolecules on a long time scale [89–93]. In this high-resolution model, the amino acid residues are represented by Cα, Cβ, the center of mass of side chains and another pseudoatom placed in the center of the Cα-Cα pseudo-bond. In this model, the amino acid residues are represented by Cα, Cβ, the center of mass of side chains and the center of the Cα-Cα pseudo-bond. The CABS-flex approach implemented as a Python 2.7 object-oriented standalone package was used in this study to allow for robust conformational sampling proven to accurately recapitulate all-atom MD simulation trajectories of proteins on a long time scale. Conformational sampling in the CABS-flex approach is conducted with the aid of Monte Carlo replica-exchange dynamics and involves local moves of individual amino acids in the protein structure and global moves of small fragments. The default settings were used in which soft native-like restraints are imposed only on pairs of residues fulfilling the following conditions : the distance between their *C_α_* atoms was smaller than 8 Å, and both residues belong to the same secondary structure elements. A total of 1000 independent CG-CABS simulations were performed for each of the systems studied. In each simulation, the total number of cycles was set to 10,000 and the number of cycles between trajectory frames was 100. MODELLER-based reconstruction of simulation trajectories to all-atom representation [94] provided by the CABS-flex package was employed to produce atomistic models of equilibrium ensembles for studied systems.

### 3.3. All-Atom Molecular Dynamics Simulations

To characterize the dynamic response of the SARS-CoV-2 RBD upon antibody binding and compare the performance of simplified and atomistic simulations, we employed a dual-method framework: (a) CG-CABS simulations with full atomic backmapping across a panel of RBD–antibody complexes, and (b) explicit-solvent all-atom molecular dynamics (MD) simulations (500 ns per system) augmented with a minimal, structurally informed glycan environment. Structural analysis and all-atom MD simulations were performed according to the protocol detailed in our recent study of RBD-antibody complexes [94]. In brief, the protonation states for all the titratable residues of the antibody and RBD proteins were assigned at pH 7.0 using Propka 3.1 software and web server [95,96]. The glycan chains were built using CHARMM-GUI Glycan Reader [97,98] at glycosylation sites N331 and N343 of RBD. NAMD 2.13-multicore-CUDA package [99] with CHARMM36m force field [100] were used in all-atom MD simulations. These simulations incorporate a minimal glycan representation at key structural sites, providing a realistic assessment of steric effects and surface accessibility without the prohibitive cost of modeling full glycan ensembles. This approach ensures that the local steric footprint and chemical environment of the RBD are accurately represented, particularly in regions where glycans may modulate antibody binding or receptor interaction. Each system was solvated with TIP3P water molecules and neutralizing 0.15 M NaCl in a periodic box that extended 10 Å beyond any protein atom in the system [101] The heavy atoms in the complex were restrained using a force constant of 1000 kJ mol^−1^ nm^−1^ to perform 1ns equilibration simulation. Long-range, non-bonded van der Waals interactions were computed using an atom-based cutoff of 12 Å, with the switching function beginning at 10 Å and reaching zero at 14 Å. The SHAKE method was used to constrain all the bonds associated with hydrogen atoms. The simulations were run using a leap-frog integrator with a 2 fs integration time step. The ShakeH algorithm in NAMD was applied for the water molecule constraints. A 310 K temperature was maintained using the Nóse-Hoover thermostat with 1.0 ps time constant and 1 atm pressure was maintained using isotropic coupling to the Parrinello-Rahman barostat [102,103]. The long-range electrostatic interactions were calculated using the particle mesh Ewald method [104] with a cut-off of 1.2 nm and a fourth-order (cubic) interpolation. The simulations were performed under an NPT ensemble with a Langevin thermostat and a Nosé–Hoover Langevin piston at 310 K and 1 atm. The damping coefficient (gamma) of the Langevin thermostat was 1/ps. In NAMD, the Nosé–Hoover Langevin piston method is a combination of the Nosé–Hoover constant pressure method [105,106] and piston fluctuation control implemented using Langevin dynamics [107]. An NPT production simulation was run on equilibrated structures for 500 ns keeping the temperature at 310 K and a constant pressure (1 atm).

### 3.4. Mutational Scanning of the RBD-Antibody Binding Interfaces

Mutational scanning analysis of the binding epitope residues for the S RBD–antibody complexes. Each binding epitope residue was systematically mutated using all substitutions, and corresponding protein stability and binding free energy changes were computed. The BeAtMuSiC approach [108–110] was employed and evaluated the impact of mutations on both the strength of interactions at the protein–protein interface and the overall stability of the complex using statistical energy functions. BeAtMuSiC is a knowledge-based statistical potential was applied to 1,000 atomistic reconstructed conformations sampled from each CG-CABS ensemble. This method quantifies mutation-induced energy changes (ΔΔG) through three physically grounded terms: (a) a Lennard-Jones-like potential for van der Waals interactions parameterized on observed atomic contact frequencies in the PDB; (b) a directional hydrogen-bonding term based on geometric and distance constraints; and (c) a solvation term modeling changes in buried surface area using statistical potentials. BeAtMuSiC identifies a residue as part of the protein–protein interface if its solvent accessibility in the complex is at least 5% lower than its solvent accessibility in the individual protein partner(s). The binding free energy of the protein–protein complex can be expressed as the difference in the folding free energy of the complex and folding free energies of the two protein binding partners. The change in the binding energy due to a mutation is

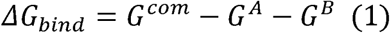

*G^eom^* is the free energy of the complex. This is the Gibbs free energy associated with the folded, bound state of the entire protein–protein complex (e.g., the Spike RBD–antibody complex). *G ^A^* is the free energy of the first binding partner (e.g., the isolated S-RBD) in its unbound, folded state. *G^B^* is the free energy of the second binding partner (e.g., the isolated antibody) in its unbound, folded state. The change in the binding energy due to a mutation was calculated then as

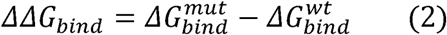

*ΔΔG_bind_* is the change in binding free energy resulting from a specific mutation. This quantifies how the mutation affects the binding affinity compared to the wild-type (original) interaction. A positive *ΔΔG_bind* typically indicates weakened binding (the mutation makes binding less favorable or more difficult), while a negative *ΔΔG_bind* indicates strengthened binding (the mutation makes binding more favorable). 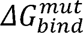 is the binding free energy calculated using Equation (4), but for the mutated protein complex (e.g., a mutant RBD bound to the antibody). 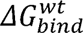 is the binding free energy calculated using Equation (4), but for the wild-type (unmutated) protein complex, serving as the reference state. We leveraged rapid calculations based on statistical potentials to compute the ensemble-averaged binding free energy changes using equilibrium samples from simulation trajectories. The binding free energy changes were obtained by averaging over 1000 and 10,000 equilibrium samples for each of the systems studied.

### 3.5. Binding Free Energy Computations of the RBD Complexes with Antibodies

MM-GBSA approach [111–117] with the AMBER21 suite [118] was employed for rigorous validation and residue-level decomposition. This dual framework leverages BeAtMuSiC efficiency for rapid mutational screening and physical rigor of MM-GBSA computations for mechanistic dissection of van der Waals and electrostatic contributions to binding and identification of binding affinity hotspots. We calculated the ensemble-averaged changes in binding free energy using 1000 equilibrium samples obtained from simulation trajectories for each system under study. Initially, the binding free energies of the RBD–antibody complexes were assessed using the MM-GBSA approach. Additionally, we conducted an energy decomposition analysis to evaluate the contribution of each amino acid during the binding of the RBD to antibodies. The binding free energy for the RBD–antibody complex was obtained using:

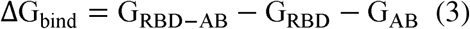

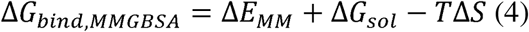

where ΔE_MM_ is total gas phase energy (sum of ΔEinternal, ΔEelectrostatic, and ΔEvdw); ΔGsol is sum of polar (ΔGGB) and non-polar (ΔGSA) contributions to solvation. Here, G _RBD–ANTIBODY_ represent the average over the snapshots of a single trajectory of the complex, G_RBD_ and G_ANTIBODY_ corresponds to the free energy of RBD and antibody respectively.

The polar and non-polar contributions to the solvation free energy were calculated using a Generalized Born solvent model and consideration of the solvent-accessible surface area. MM-GBSA was employed to predict the binding free energy and decompose the free energy contributions to the binding free energy of a protein–protein complex on a per residue basis. The binding free energy with MM-GBSA was computed by averaging the results of computations over 10,000 samples from the equilibrium ensembles. The standard error of the mean (SEM) for binding free energy estimates was calculated from the distribution of values obtained across the 10,000 snapshots sampled for each system. In this study, we chose the “single trajectory” protocol (one trajectory of the complex) because it is less noisy due to the cancellation of intermolecular energy contributions. Entropy calculations typically dominate the computational cost of MM-GBSA estimates. In this study, the entropy contribution was not included in the calculations of binding free energies of the RBD–antibody complexes because the entropic differences in estimates of relative binding affinities were expected to be small owing to the small mutational changes and the preservation of the conformational dynamics. MM-GBSA energies were evaluated with the MMPBSA.py script in the AmberTools21 package [119] and gmx_MMPBSA, a new tool to perform end-state free energy calculations from CHARMM and GROMACS trajectories [120].

### 3.6. Local Frustration Analysis of Conformational Ensembles

To characterize the energetically encoded signatures of functional sites, we employed a local frustration analysis framework rooted in energy landscape theory [68–70]. This approach quantifies the extent to which local interactions within the native protein structure are energetically optimized or strained, distinguishing between residues that are evolutionarily and structurally stabilized from those involved in conformational adaptability or functional stress. Two complementary frustration indices were computed. Configurational (or conformational) frustration assesses the sensitivity of a native contact to local structural perturbations. It is calculated by randomizing both residue identities and interatomic distances within the native contact geometry, thereby measuring whether the observed interaction geometry is more favorable than structurally plausible alternatives. Mutational frustration evaluates the energetic optimality of a given residue–residue pair by comparing its interaction energy to that of all possible amino acid substitutions at the same positions, while the backbone conformation remains fixed. This metric captures evolutionary constraints by reflecting whether the native amino acid pairing is more favorable than evolutionary accessible alternatives [68,69]. Both frustration indices were expressed as Z-scores, defined as:

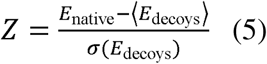

where *E*_native_is the energy of the native contact, and (*E*_decoys_) and a-(*E*_decoys_)represent the mean and standard deviation, respectively, of energies computed from an ensemble of 1,000 structural decoys generated by perturbing local conformations or residue identities.

Following well-established thresholds validated across diverse protein systems^68,69^, contacts were classified as minimally frustrated (Z > 0.78). In this regime, the native interaction is significantly more favorable than decoy alternatives, indicating strong energetic optimization. Highly frustrated (Z < −1.0). In the high frustration regime, the native interaction is significantly less favorable, reflecting local strain or conflict. Neutrally frustrated (−1.0 ≤ Z ≤ 0.78) which is a regime when the native interaction is neither strongly favored nor disfavored, consistent with conformational plasticity or mutational tolerance. To map these interactions onto a residue-level profile, the local density of frustrated contacts within a 5 Å radius of each residue was computed. A residue was assigned a dominant frustration category if more than 50% of its interacting contacts within the most populated conformational ensemble belonged to the same frustration class.

## 5. Conclusions

By integrating conformational dynamics, mutational scanning, MM-GBSA binding energetics, and frustration profiling, this study has dissected the molecular mechanisms of three classes of broadly neutralizing XGI antibodies targeting the super-conserved SARS-CoV-2 RBD epitopes SCORE-A, SCORE-B, and SCORE-C. Our results establish a unifying biophysical framework that explains the experimentally observed gradient of neutralization potency and escape resistance across these classes. Across all three epitopes, residues that repeatedly mutate in circulating variants—such as K356, V503, Y508, S383, and T385—consistently reside in zones of neutral frustration, energetically permissive positions that tolerate substitutions without destabilizing the RBD. These neutral-frustration “playgrounds” provide the virus with low-cost escape routes, explaining why antibodies that rely on such hotspots, like the SCORE-A antibody XGI-183, are vulnerable to immune evasion. In contrast, antibodies that anchor their binding to minimally frustrated, evolutionarily constrained cores—notably the SCORE-C antibody XGI-171, and to a lesser extent the SCORE-B antibody XGI-198—create a high genetic barrier to resistance. The relationship between conformational and mutational frustration distributions encodes the distinct mechanisms: aligned neutral frustration yields permissive, plastic interfaces (SCORE-A); subtle decoupling improves escape resistance while retaining the scaffold (XGI-198); pronounced decoupling enables targeting of constrained cores with maintained adaptability (XGI-203); and convergence of minimal frustration in both distributions creates an invulnerable, allosteric interface (XGI-171). Broad neutralization thus arises not from ultra-high-affinity anchors but from strategic distribution of binding energy across multiple minimally frustrated contacts, a “distributed redundancy” model that explains why XGI-171 binds all tested variants yet neutralizes weakly, and why XGI-198 outranks XGI-203 within the same SCORE class. Our integrated computational framework provides a predictive roadmap for antibody and vaccine design: frustration profiling can prospectively rank epitopes by their content of minimally frustrated residues, and immunogens that focus the immune response on evolutionarily constrained cores may elicit antibodies that the virus cannot easily evade.

## Author Contributions

Conceptualization, G.V.; Methodology, M.A., W.G., M.L., L.T., B.F., G.V.; Software, W.G., M.L., L.T., B.F., G.V.; Validation, W.G., M.L., L.T., B.F., G.V.; Formal analysis, W.G., M.L., L.T., B.F., G.V.; Resources, W.G., M.L., L.T., B.F., G.V.; Data curation, W.G., M.L., L.T., B.F., G.V.; Writing-original draft preparation, G.V.; Writing-review and editing, G.V.; Visualization. W.G., M.L., L.T., B.F., G.V.; Supervision G.V.; Project administration, G.V.;. Funding acquisition, G.V. All authors have read and agreed to the published version of the manuscript.

## Data Availability Statement

Data is fully contained within the article and Supplementary Materials. Crystal structures were obtained and downloaded from the Protein Data Bank (http://www.rcsb.org). The rendering of protein structures was done with UCSF ChimeraX package (https://www.rbvi.ucsf.edu/chimerax/) and Pymol (https://pymol.org/2/). All mutational heatmaps were produced using the developed software that is freely available at https://alshahrani.shinyapps.io/HeatMapViewerApp/.

## Acknowledgments

The authors acknowledge support from Schmid College of Science and Technology at Chapman University for providing computing resources at the Keck Center for Science and Engineering.

