## Supplemental Figures S1-S8, Tables S1-S9 for "Frustration Landscapes of Broadly Neutralizing SARS-CoV-2 Spike Antibodies Targeting Conserved Epitopes Reveal Energetic Logic of Escape-Proof and Escape-Prone Mechanisms"

**Supp****lementary Material**

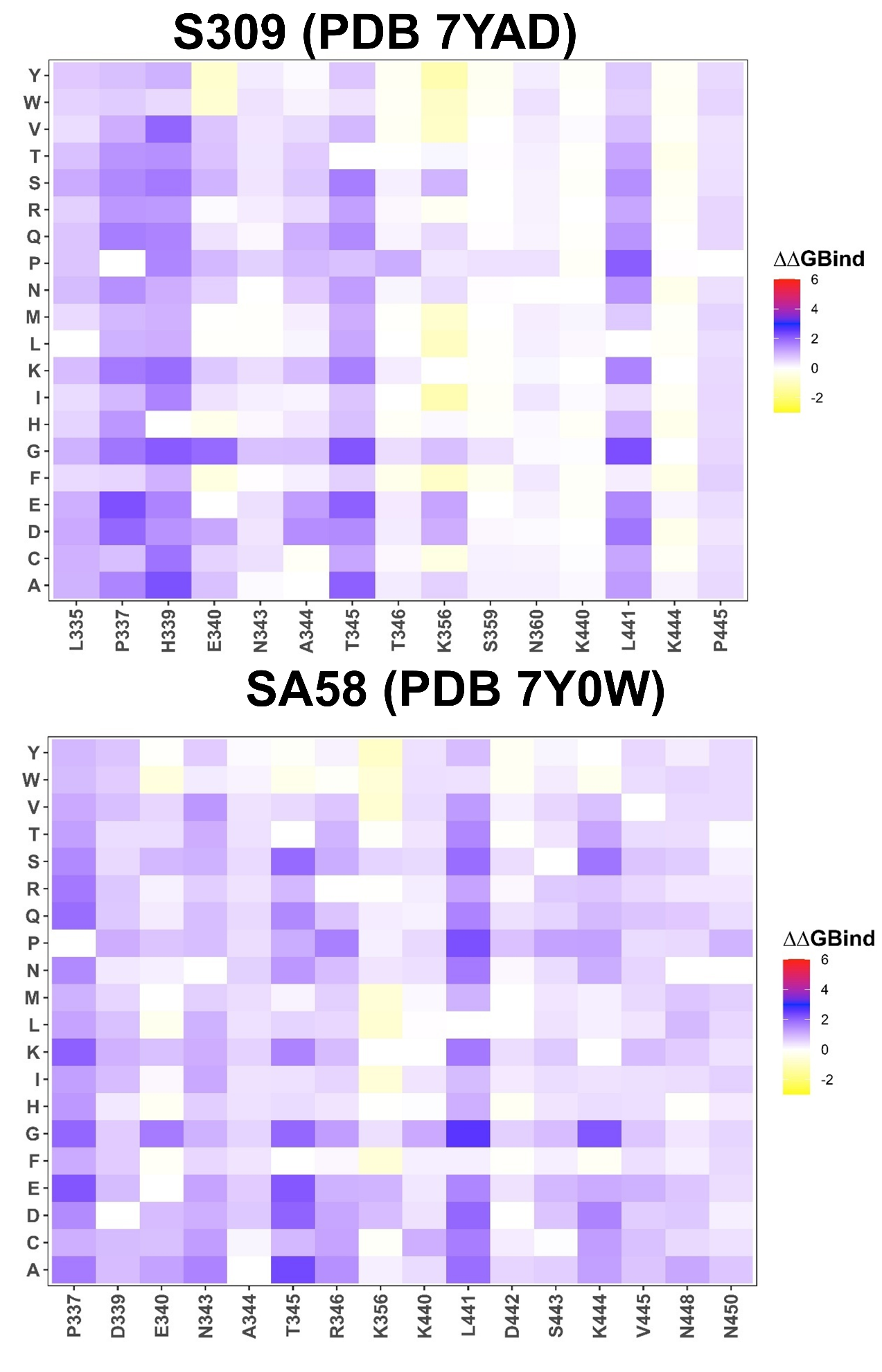

**Figure S1. Mutational scanning of binding for the RBD complexes with SCORE-A epitope class S309 and SA58 antibodies.** The mutational scanning heatmaps for the binding epitope residues in the S-RBD complexes with S309 (top panel) and SA58 (bottom panel). The heatmaps show the computed binding free energy changes for 20 single mutations on the sites of variants. The squares on the heatmap are colored using a 4-colored scale blue-white-yellow-red, with blue indicating the largest unfavorable effect on binding and stability, while yellow-red points to mutations that have favorable effect and improve binding.

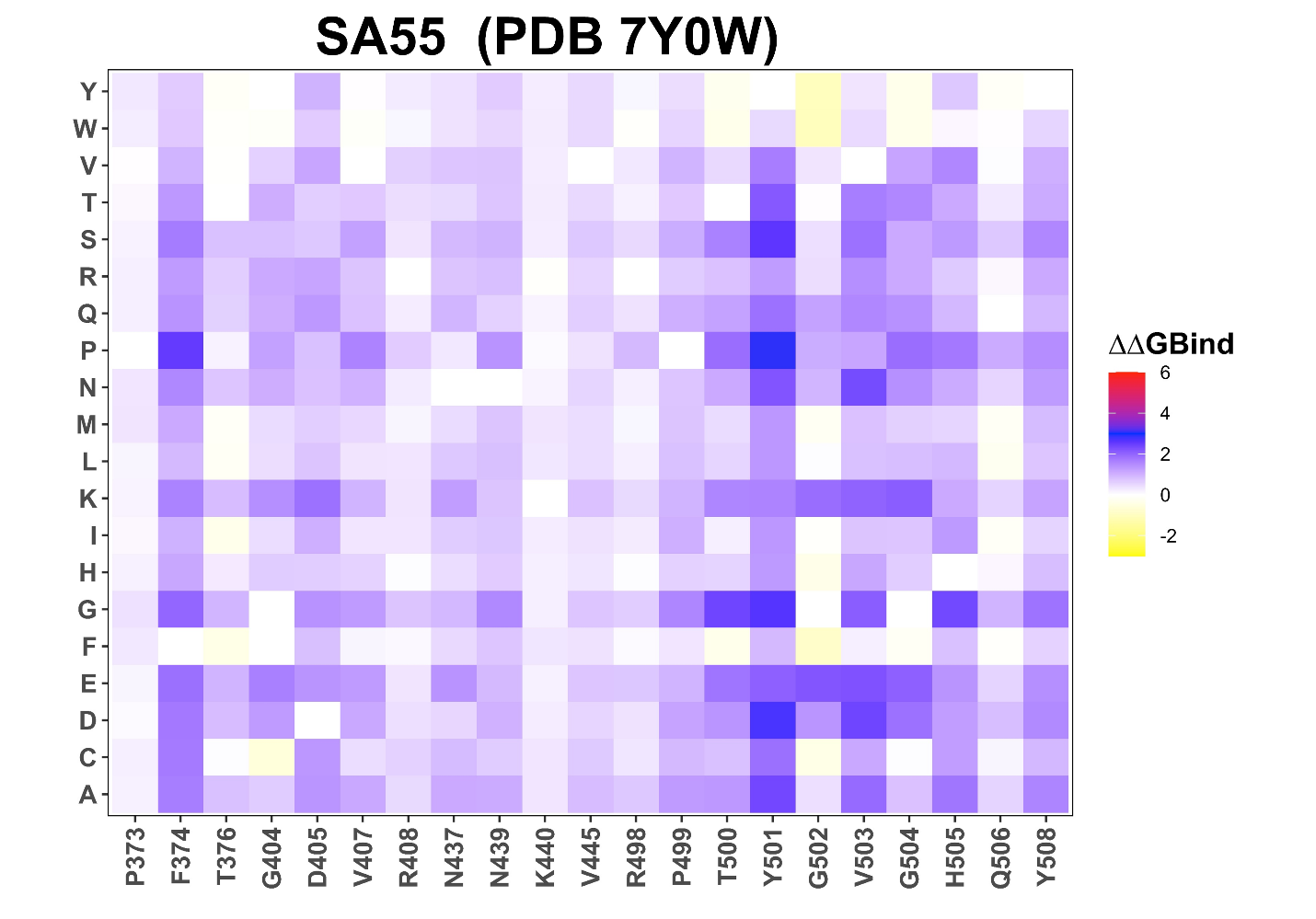

**Figure S2. Mutational scanning of binding for the RBD complexes with SCORE-B epitope class SA55 antibody.** The mutational scanning heatmaps for the binding epitope residues in the S-RBD complex with SA55 antibody. The heatmaps show the computed binding free energy changes for 20 single mutations on the sites of variants. The squares on the heatmap are colored using a 4-colored scale blue-white-yellow-red, with blue indicating the largest unfavorable effect on binding and stability, while yellow-red points to mutations that have favorable effect and improve binding.

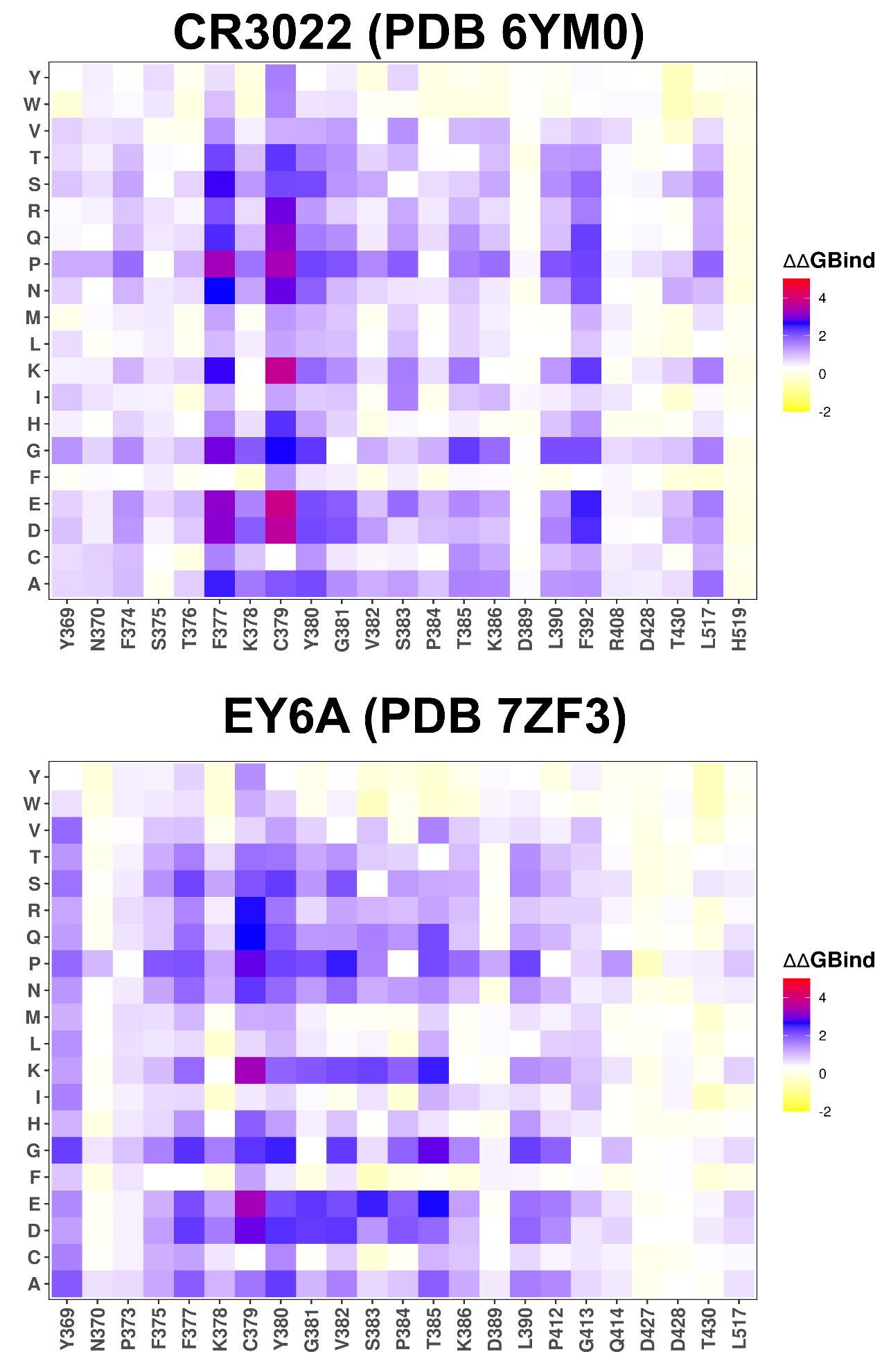

**Figure S3. Mutational scanning of binding for the RBD complexes with SCORE-C epitope class CR3022 and EY6A antibodies.** The mutational scanning heatmaps for the binding epitope residues in the S-RBD complexes with CR3022 (top panel) and EY6A (bottom panel). The heatmaps show the computed binding free energy changes for 20 single mutations on the sites of variants. The squares on the heatmap are colored using a 4-colored scale blue-white-yellow-red, with blue indicating the largest unfavorable effect on binding and stability, while yellow-red points to mutations that have favorable effect and improve binding.

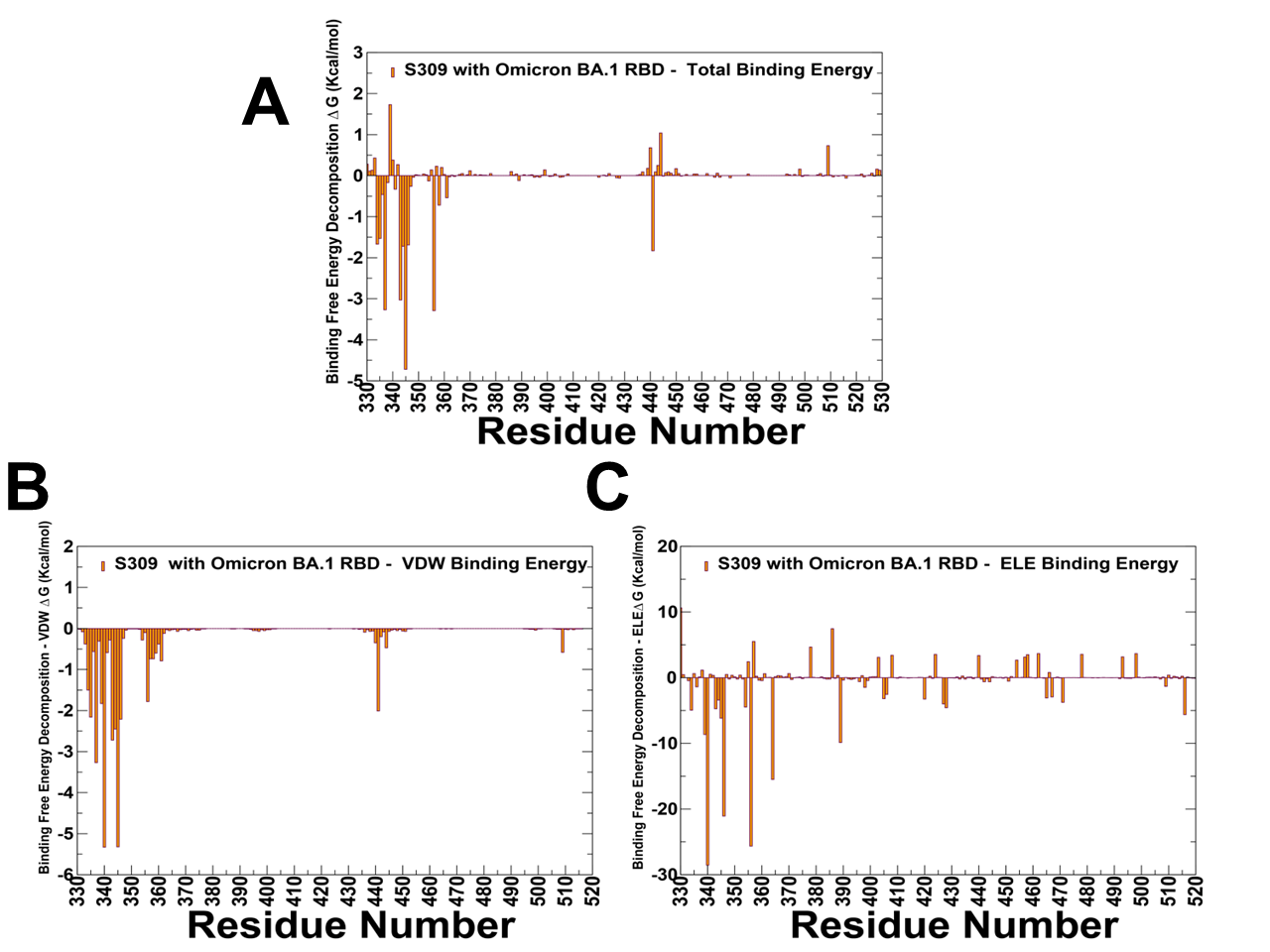

**Figure S4.** The residue-based decomposition of the binding MM-GBSA energies (A), van der Waals contributions (B) and electrostatic interactions (C) for the S-RBD complexes with SCORE-A S309 antibody. The binding free energy with MM-GBSA was computed by averaging the results of computations over 10,000 samples from the equilibrium ensembles. The standard error of the mean (SEM) for binding free energy estimates was calculated from the distribution of values obtained across the 10,000 snapshots sampled for each system. The statistical errors was estimated on the basis of the deviation between block average and are within 0.0.9 kcal/mol.

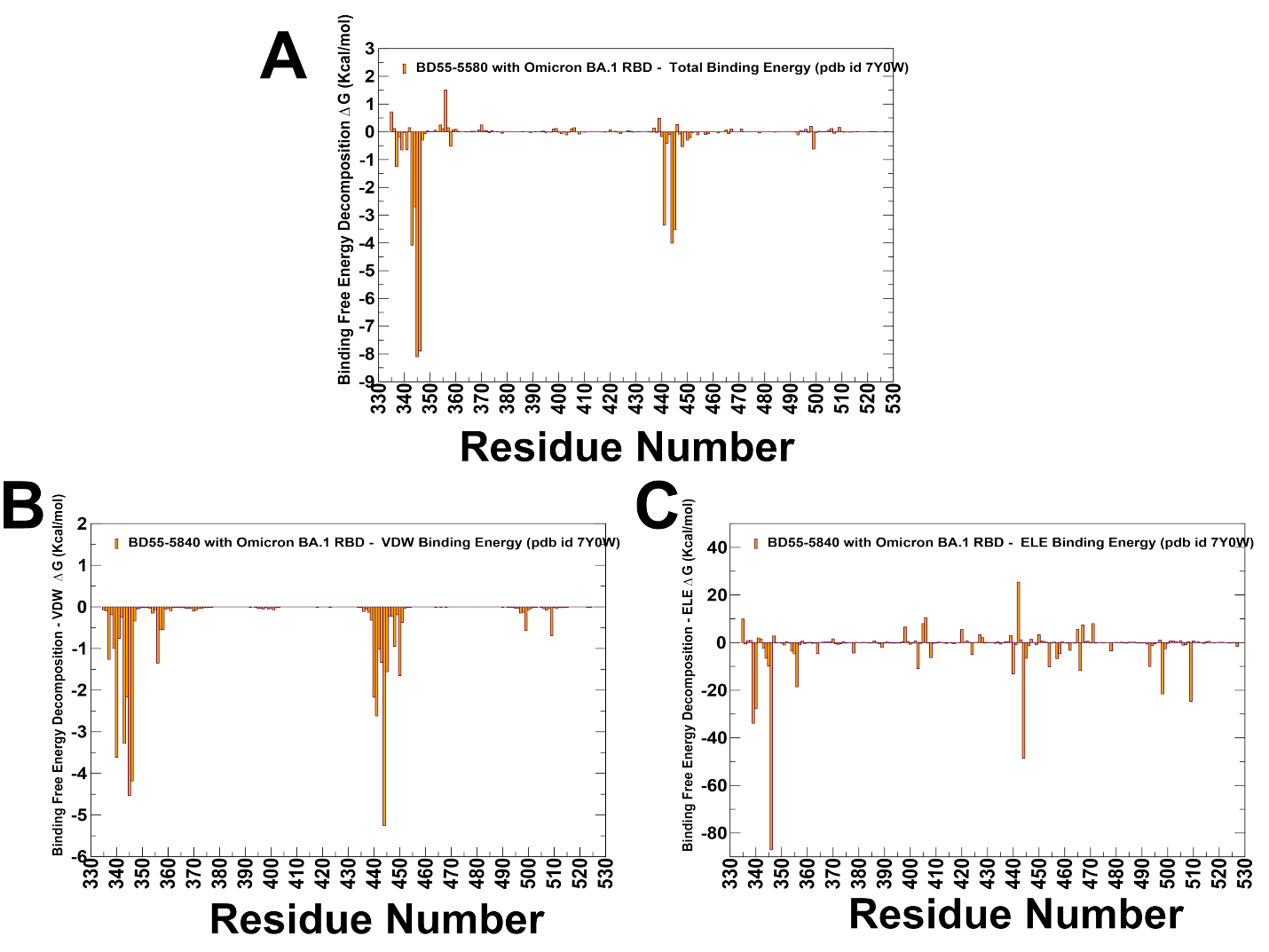

**Figure S5.** The residue-based decomposition of the binding MM-GBSA energies (A), van der Waals contributions (B) and electrostatic interactions (C) for the S-RBD complexes with SCORE-A SA58 antibody. The binding free energy with MM-GBSA was computed by averaging the results of computations over 10,000 samples from the equilibrium ensembles. The standard error of the mean (SEM) for binding free energy estimates was calculated from the distribution of values obtained across the 10,000 snapshots sampled for each system. The statistical errors was estimated on the basis of the deviation between block average and are within 0.15 kcal/mol.

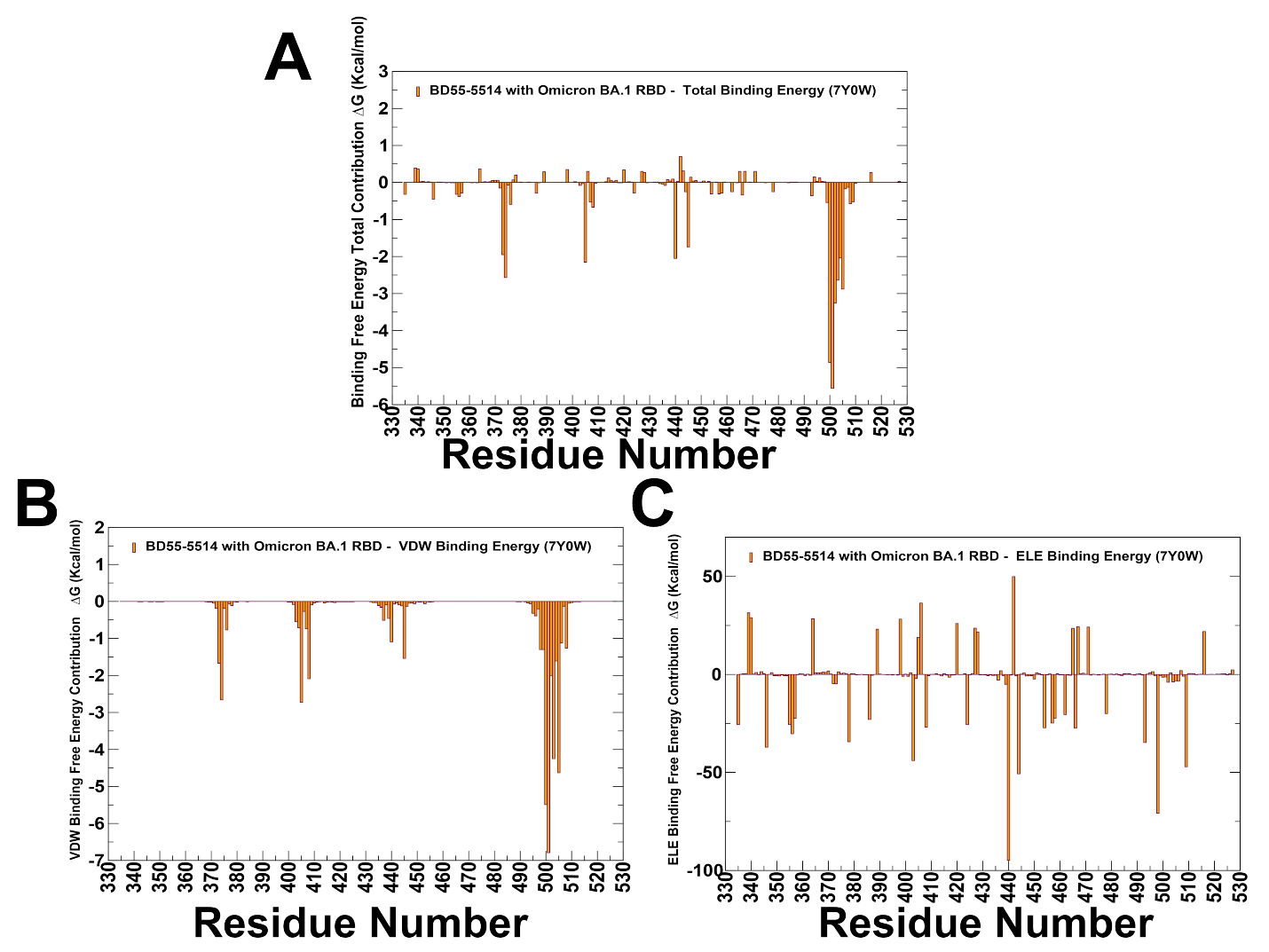

**Figure S6.** The residue-based decomposition of the binding MM-GBSA energies (A), van der Waals contributions (B) and electrostatic interactions (C) for the S-RBD complexes with SCORE-B SA55 antibody. The binding free energy with MM-GBSA was computed by averaging the results of computations over 10,000 samples from the equilibrium ensembles. The standard error of the mean (SEM) for binding free energy estimates was calculated from the distribution of values obtained across the 10,000 snapshots sampled for each system. The statistical errors was estimated on the basis of the deviation between block average and are within 0.22 kcal/mol.

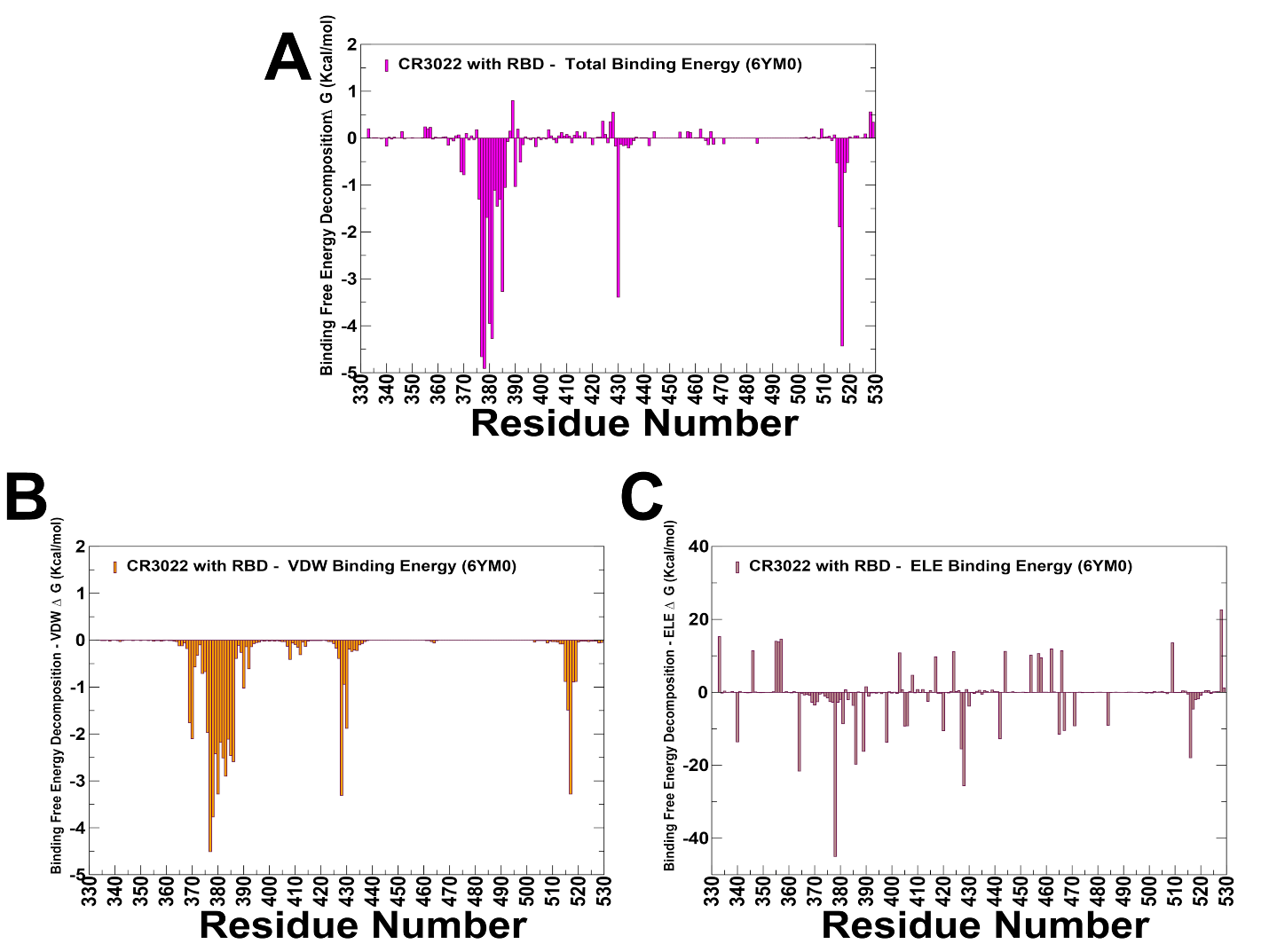

**Figure S7.** The residue-based decomposition of the binding MM-GBSA energies (A), van der Waals contributions (B) and electrostatic interactions (C) for the S-RBD complexes with SCORE-C CR3022 antibody. The binding free energy with MM-GBSA was computed by averaging the results of computations over 10,000 samples from the equilibrium ensembles. The standard error of the mean (SEM) for binding free energy estimates was calculated from the distribution of values obtained across the 10,000 snapshots sampled for each system. The statistical errors was estimated on the basis of the deviation between block average and are within 0.11 kcal/mol.

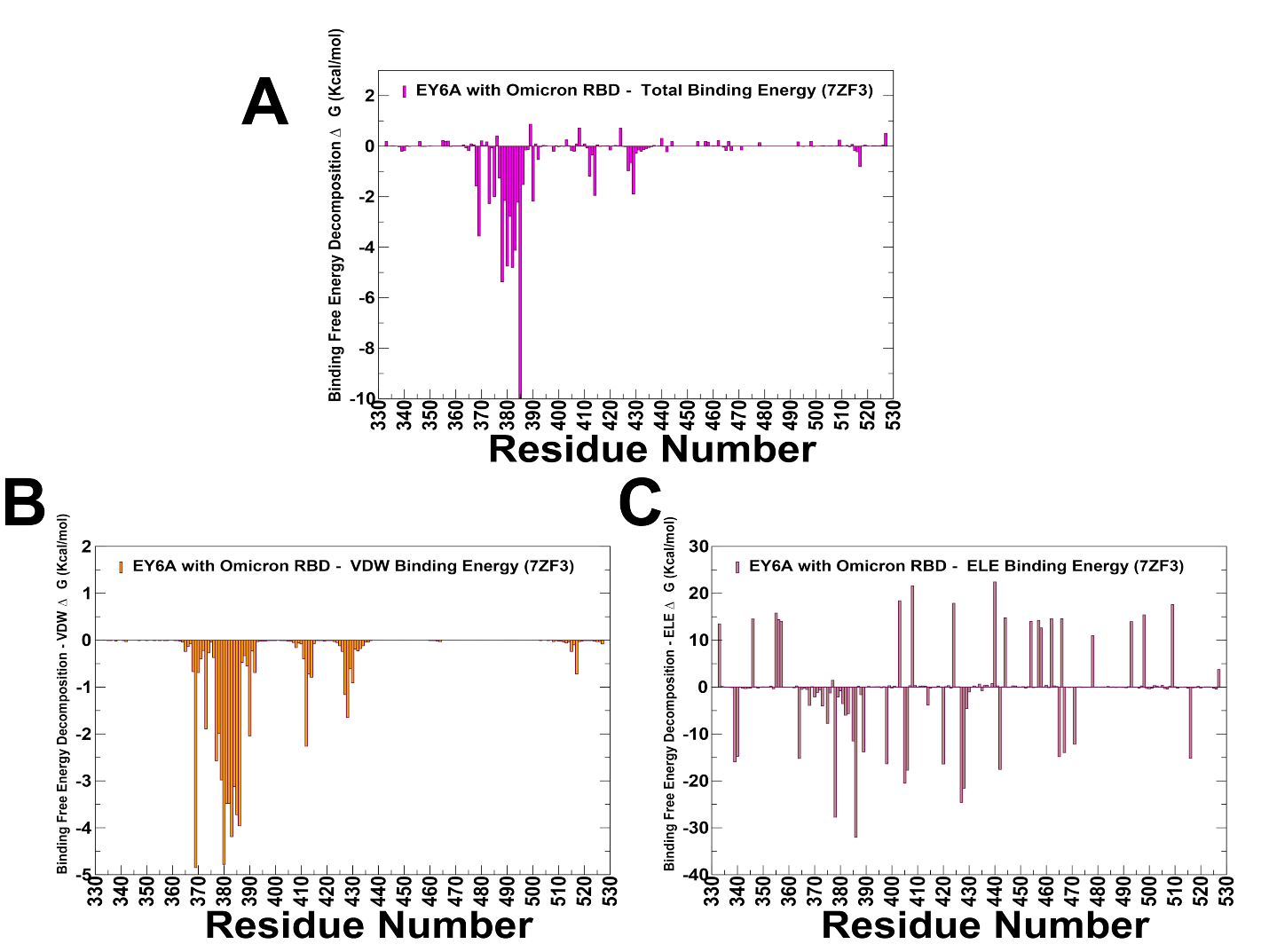

**Figure S8.** The residue-based decomposition of the binding MM-GBSA energies (A), van der Waals contributions (B) and electrostatic interactions (C) for the S-RBD complexes with SCORE-C EY6A antibody. The binding free energy with MM-GBSA was computed by averaging the results of computations over 10,000 samples from the equilibrium ensembles. The standard error of the mean (SEM) for binding free energy estimates was calculated from the distribution of values obtained across the 10,000 snapshots sampled for each system. The statistical errors was estimated on the basis of the deviation between block average and are within 0.15-0.17 kcal/mol.

**Table S1. The list of the intermolecular contacts in the structure of the XGI-183 antibody complex with RBD (pdb id 9KZE).** The interfacial contacts in the structure are defined by counting the number of interatomic contacts within a 5.5 Å distance threshold between atoms of the interacting proteins.

| **RBD Residue** | **RBD Residue Number** | **RBD chain** | **Ab Residue** | **Ab Residue Number** | **Ab chain** |
| --- | --- | --- | --- | --- | --- |
| PRO | 337 | A | GLY | 29 | L |
| PRO | 337 | A | ASN | 80 | L |
| GLU | 340 | A | GLY | 29 | L |
| GLU | 340 | A | SER | 36 | L |
| GLU | 340 | A | ASN | 80 | L |
| GLU | 340 | A | SER | 83 | L |
| GLU | 340 | A | GLY | 84 | L |
| VAL | 341 | A | GLY | 29 | L |
| VAL | 341 | A | SER | 36 | L |
| ASN | 343 | A | ASN | 27 | L |
| ALA | 344 | A | ASN | 27 | L |
| ALA | 344 | A | SER | 36 | L |
| THR | 345 | A | ASN | 26 | L |
| THR | 345 | A | ASN | 27 | L |
| THR | 346 | A | ASN | 26 | L |
| THR | 346 | A | ASN | 27 | L |
| THR | 346 | A | SER | 109 | L |
| PHE | 347 | A | SER | 109 | L |
| ALA | 348 | A | SER | 109 | L |
| SER | 349 | A | SER | 109 | L |
| TYR | 351 | A | TRP | 107 | L |
| TYR | 351 | A | SER | 109 | L |
| TYR | 351 | A | SER | 110 | L |
| TYR | 351 | A | PHE | 113 | L |
| TYR | 351 | A | ASP | 114 | L |
| ALA | 352 | A | TRP | 107 | L |
| ALA | 352 | A | ASP | 108 | L |
| ALA | 352 | A | SER | 109 | L |
| ALA | 352 | A | SER | 110 | L |
| ALA | 352 | A | ASP | 114 | L |
| TRP | 353 | A | LEU | 110 | H |
| TRP | 353 | A | TRP | 107 | L |
| ASN | 354 | A | SER | 36 | L |
| ASN | 354 | A | LYS | 37 | L |
| ASN | 354 | A | TRP | 107 | L |
| ASN | 354 | A | ASP | 108 | L |
| ASN | 354 | A | SER | 109 | L |
| ARG | 355 | A | LEU | 110 | H |
| ARG | 355 | A | THR | 112 | H |
| ARG | 355 | A | ASN | 38 | L |
| ARG | 355 | A | TRP | 107 | L |
| LYS | 356 | A | ILE | 28 | L |
| LYS | 356 | A | GLY | 29 | L |
| LYS | 356 | A | SER | 36 | L |
| LYS | 356 | A | LYS | 37 | L |
| LYS | 356 | A | ASN | 38 | L |
| LYS | 356 | A | VAL | 39 | L |
| LYS | 356 | A | ASP | 57 | L |
| LYS | 356 | A | ASN | 80 | L |
| ARG | 357 | A | THR | 112 | H |
| ARG | 357 | A | ASN | 38 | L |
| ARG | 357 | A | TYR | 55 | L |
| ARG | 357 | A | ASP | 56 | L |
| ARG | 357 | A | SER | 65 | L |
| ARG | 357 | A | ASP | 66 | L |
| ILE | 358 | A | ASN | 38 | L |
| SER | 359 | A | ASP | 56 | L |
| SER | 359 | A | SER | 65 | L |
| SER | 359 | A | ASP | 66 | L |
| ASN | 360 | A | SER | 65 | L |
| ASN | 360 | A | ASP | 66 | L |
| ARG | 457 | A | SER | 63 | H |
| ARG | 457 | A | ASN | 64 | H |
| LEU | 461 | A | ASN | 64 | H |
| LYS | 462 | A | TYR | 58 | H |
| PHE | 464 | A | LEU | 110 | H |
| GLU | 465 | A | SER | 57 | H |
| GLU | 465 | A | TYR | 58 | H |
| GLU | 465 | A | ASN | 64 | H |
| GLU | 465 | A | LEU | 110 | H |
| ARG | 466 | A | ASN | 64 | H |
| ARG | 466 | A | GLU | 107 | H |
| ARG | 466 | A | LEU | 110 | H |
| ARG | 466 | A | THR | 112 | H |
| ARG | 466 | A | TRP | 107 | L |
| ARG | 466 | A | TRP | 116 | L |
| ASP | 467 | A | ASN | 64 | H |
| ILE | 468 | A | VAL | 55 | H |
| ILE | 468 | A | ASN | 64 | H |
| ILE | 468 | A | LYS | 65 | H |
| ILE | 468 | A | HIS | 66 | H |
| ILE | 468 | A | TRP | 107 | L |
| ILE | 468 | A | PHE | 113 | L |
| ILE | 468 | A | ASP | 114 | L |
| ILE | 468 | A | TRP | 116 | L |
| SER | 469 | A | SER | 63 | H |
| SER | 469 | A | ASN | 64 | H |
| SER | 469 | A | LYS | 65 | H |
| SER | 469 | A | HIS | 66 | H |
| THR | 470 | A | LYS | 65 | H |
| THR | 470 | A | HIS | 66 | H |
| THR | 470 | A | LYS | 72 | H |
| THR | 470 | A | PHE | 113 | L |
| GLU | 471 | A | LYS | 65 | H |
| GLU | 471 | A | HIS | 66 | H |
| GLU | 471 | A | TYR | 67 | H |
| GLU | 471 | A | LYS | 72 | H |
| LEU | 492 | A | PHE | 113 | L |

**Table S2. The list of the intermolecular contacts in the structure of the S309 antibody complex with RBD (pdb id 7YAD).** The interfacial contacts in the structure are defined by counting the number of interatomic contacts within a 5.5 Å distance threshold between atoms of the interacting proteins.

| **RBD Residue** | **RBD Residue Number** | **RBD chain** | **Ab Residue** | **Ab Residue Number** | **Ab chain** |
| --- | --- | --- | --- | --- | --- |
| ASN | 334 | M | TYR | 54 | A |
| ASN | 334 | M | THR | 30 | A |
| ASN | 334 | M | TRP | 105 | A |
| LEU | 335 | M | SER | 31 | A |
| LEU | 335 | M | PRO | 28 | A |
| LEU | 335 | M | TRP | 105 | A |
| CYS | 336 | M | TRP | 105 | A |
| CYS | 336 | M | SER | 31 | A |
| PRO | 337 | M | PHE | 106 | A |
| PRO | 337 | M | TRP | 105 | A |
| ASP | 339 | M | TYR | 32 | A |
| ASP | 339 | M | LEU | 110 | A |
| ASP | 339 | M | TYR | 100 | A |
| ASP | 339 | M | SER | 31 | A |
| GLU | 340 | M | GLY | 107 | A |
| GLU | 340 | M | TRP | 105 | A |
| GLU | 340 | M | ARG | 102 | A |
| GLU | 340 | M | GLY | 103 | A |
| GLU | 340 | M | SER | 31 | A |
| GLU | 340 | M | PHE | 106 | A |
| GLU | 340 | M | ALA | 104 | A |
| GLU | 340 | M | GLU | 108 | A |
| GLU | 340 | M | LEU | 110 | A |
| VAL | 341 | M | LEU | 110 | A |
| VAL | 341 | M | PHE | 106 | A |
| ASN | 343 | M | TYR | 100 | A |
| ASN | 343 | M | LEU | 110 | A |
| ASN | 343 | M | ILE | 111 | A |
| ASN | 343 | M | SER | 109 | A |
| ALA | 344 | M | SER | 109 | A |
| ALA | 344 | M | LEU | 110 | A |
| ALA | 344 | M | GLU | 108 | A |
| ALA | 344 | M | ILE | 111 | A |
| THR | 345 | M | LEU | 110 | A |
| THR | 345 | M | THR | 32 | B |
| THR | 345 | M | ILE | 111 | A |
| THR | 345 | M | SER | 33 | B |
| THR | 345 | M | HIS | 92 | B |
| THR | 345 | M | SER | 109 | A |
| ARG | 346 | M | GLU | 108 | A |
| ARG | 346 | M | SER | 109 | A |
| ARG | 346 | M | SER | 30 | B |
| ARG | 346 | M | ASP | 93 | B |
| ASN | 354 | M | GLU | 108 | A |
| LYS | 356 | M | PHE | 106 | A |
| LYS | 356 | M | GLU | 108 | A |
| ARG | 357 | M | PHE | 106 | A |
| ILE | 358 | M | TRP | 105 | A |
| ILE | 358 | M | PHE | 106 | A |
| SER | 359 | M | TRP | 105 | A |
| SER | 359 | M | TYR | 54 | A |
| ASN | 360 | M | TRP | 105 | A |
| CYS | 361 | M | TRP | 105 | A |
| LYS | 440 | M | SER | 31 | B |
| LEU | 441 | M | SER | 31 | B |
| LEU | 441 | M | THR | 32 | B |
| LEU | 441 | M | ILE | 111 | A |
| ARG | 509 | M | THR | 32 | B |
| ARG | 509 | M | ILE | 111 | A |

**Table S3. The list of the intermolecular contacts in the structure of the SA58 antibody complex with RBD (pdb id 7Y0W).** The interfacial contacts in the structure are defined by counting the number of interatomic contacts within a 5.5 Å distance threshold between atoms of the interacting proteins.

| **RBD Residue** | **RBD Residue Number** | **RBD chain** | **Ab Residue** | **Ab Residue Number** | **Ab chain** |
| --- | --- | --- | --- | --- | --- |
| PRO | 337 | R | LEU | 29 | L |
| PRO | 337 | R | SER | 28 | L |
| ASP | 339 | R | ASN | 95 | L |
| ASP | 339 | R | GLU | 1 | L |
| GLU | 340 | R | ALA | 27 | L |
| GLU | 340 | R | SER | 28 | L |
| GLU | 340 | R | ASN | 95 | L |
| GLU | 340 | R | GLU | 1 | L |
| GLU | 340 | R | ARG | 26 | L |
| GLU | 340 | R | GLY | 30 | L |
| GLU | 340 | R | LEU | 29 | L |
| GLU | 340 | R | VAL | 2 | L |
| VAL | 341 | R | ASN | 95 | L |
| VAL | 341 | R | LEU | 29 | L |
| ASN | 343 | R | SER | 94 | L |
| ASN | 343 | R | PRO | 97 | L |
| ASN | 343 | R | ASN | 95 | L |
| ASN | 343 | R | TRP | 96 | L |
| ALA | 344 | R | SER | 94 | L |
| ALA | 344 | R | ASN | 95 | L |
| ALA | 344 | R | TRP | 96 | L |
| THR | 345 | R | TRP | 96 | L |
| THR | 345 | R | TYR | 105 | H |
| THR | 345 | R | ASP | 34 | L |
| THR | 345 | R | TYR | 93 | L |
| THR | 345 | R | LEU | 98 | L |
| THR | 345 | R | SER | 94 | L |
| THR | 345 | R | ASN | 95 | L |
| ARG | 346 | R | PHE | 106 | H |
| ARG | 346 | R | SER | 94 | L |
| ARG | 346 | R | SER | 103 | H |
| ARG | 346 | R | ASP | 104 | H |
| ARG | 346 | R | TYR | 105 | H |
| ARG | 346 | R | ASP | 34 | L |
| ARG | 346 | R | TYR | 93 | L |
| LYS | 356 | R | LEU | 29 | L |
| ARG | 357 | R | LEU | 29 | L |
| ILE | 358 | R | LEU | 29 | L |
| LYS | 440 | R | TRP | 50 | H |
| LYS | 440 | R | THR | 57 | H |
| LYS | 440 | R | ASN | 32 | H |
| LYS | 440 | R | PRO | 58 | H |
| LYS | 440 | R | ASN | 52 | H |
| LYS | 440 | R | TYR | 102 | H |
| LYS | 440 | R | THR | 59 | H |
| LEU | 441 | R | ASN | 52 | H |
| LEU | 441 | R | TRP | 96 | L |
| LEU | 441 | R | TYR | 102 | H |
| LEU | 441 | R | TRP | 50 | H |
| LEU | 441 | R | SER | 103 | H |
| LEU | 441 | R | TYR | 105 | H |
| ASP | 442 | R | TYR | 102 | H |
| ASP | 442 | R | SER | 103 | H |
| ASP | 442 | R | TYR | 105 | H |
| SER | 443 | R | ASP | 54 | H |
| SER | 443 | R | TYR | 102 | H |
| SER | 443 | R | ASN | 32 | H |
| LYS | 444 | R | THR | 30 | H |
| LYS | 444 | R | ASN | 32 | H |
| LYS | 444 | R | SER | 31 | H |
| LYS | 444 | R | ASP | 54 | H |
| LYS | 444 | R | TYR | 102 | H |
| VAL | 445 | R | ASP | 54 | H |
| ASN | 448 | R | TYR | 102 | H |
| ASN | 448 | R | SER | 103 | H |
| ASN | 450 | R | TYR | 102 | H |
| ASN | 450 | R | SER | 103 | H |
| TYR | 451 | R | SER | 103 | H |
| ARG | 509 | R | TYR | 105 | H |
| ARG | 509 | R | TRP | 96 | L |

**Table S4. The list of the intermolecular contacts in the structure of the XGI-188 antibody complex with RBD (pdb id 9L05).** The interfacial contacts in the structure are defined by counting the number of interatomic contacts within a 5.5 Å distance threshold between atoms of the interacting proteins.

| **RBD Residue** | **RBD Residue Number** | **RBD chain** | **Ab Residue** | **Ab Residue Number** | **Ab chain** |
| --- | --- | --- | --- | --- | --- |
| PRO | 373 | A | SER | 59 | H |
| PHE | 375 | A | SER | 59 | H |
| ASN | 437 | A | HIS | 57 | H |
| ASN | 437 | A | SER | 59 | H |
| ASN | 437 | A | ASN | 64 | H |
| SER | 438 | A | ASN | 64 | H |
| ASN | 439 | A | TYR | 38 | H |
| ASN | 439 | A | ASN | 64 | H |
| ASN | 439 | A | TYR | 66 | H |
| ASN | 439 | A | ASN | 114 | L |
| LYS | 440 | A | ASN | 64 | H |
| LYS | 440 | A | THR | 65 | H |
| LYS | 440 | A | TYR | 66 | H |
| LYS | 444 | A | ASN | 109 | L |
| PRO | 445 | A | ASN | 109 | L |
| PRO | 445 | A | SER | 113 | L |
| ARG | 498 | A | LEU | 112 | H |
| ARG | 498 | A | ASP | 36 | L |
| ARG | 498 | A | ASN | 37 | L |
| ARG | 498 | A | ASN | 109 | L |
| PRO | 499 | A | TYR | 38 | H |
| PRO | 499 | A | TYR | 66 | H |
| PRO | 499 | A | VAL | 112 | H |
| PRO | 499 | A | ASN | 109 | L |
| PRO | 499 | A | SER | 113 | L |
| PRO | 499 | A | ASN | 114 | L |
| THR | 500 | A | VAL | 109 | H |
| THR | 500 | A | VAL | 112 | H |
| THR | 500 | A | LEU | 112 | H |
| THR | 500 | A | ASP | 36 | L |
| THR | 500 | A | ASN | 37 | L |
| THR | 500 | A | TYR | 107 | L |
| THR | 500 | A | ASP | 108 | L |
| THR | 500 | A | ASN | 109 | L |
| THR | 500 | A | SER | 113 | L |
| THR | 500 | A | ASN | 114 | L |
| TYR | 501 | A | TYR | 38 | H |
| TYR | 501 | A | VAL | 109 | H |
| TYR | 501 | A | VAL | 112 | H |
| TYR | 501 | A | LEU | 112 | H |
| GLY | 502 | A | VAL | 109 | H |
| GLY | 502 | A | GLY | 110 | H |
| GLY | 502 | A | GLY | 111 | H |
| GLY | 502 | A | VAL | 112 | H |
| GLY | 502 | A | LEU | 112 | H |
| VAL | 503 | A | GLU | 35 | H |
| VAL | 503 | A | TYR | 36 | H |
| VAL | 503 | A | TYR | 38 | H |
| VAL | 503 | A | HIS | 57 | H |
| VAL | 503 | A | VAL | 109 | H |
| VAL | 503 | A | GLY | 110 | H |
| GLY | 504 | A | TYR | 36 | H |
| GLY | 504 | A | VAL | 109 | H |
| GLN | 506 | A | TYR | 38 | H |
| GLN | 506 | A | ASN | 64 | H |
| GLN | 506 | A | TYR | 66 | H |
| GLN | 506 | A | VAL | 109 | H |
| TYR | 508 | A | HIS | 57 | H |
| TYR | 508 | A | SER | 59 | H |

**Table S5. The list of the intermolecular contacts in the structure of the XGI-203 antibody complex with RBD (pdb id 9L07).** The interfacial contacts in the structure are defined by counting the number of interatomic contacts within a 5.5 Å distance threshold between atoms of the interacting proteins.

| **RBD Residue** | **RBD Residue Number** | **RBD chain** | **Ab Residue** | **Ab Residue Number** | **Ab chain** |
| --- | --- | --- | --- | --- | --- |
| ALA | 372 | A | SER | 59 | H |
| ALA | 372 | A | GLY | 63 | H |
| PRO | 373 | A | TYR | 58 | H |
| PRO | 373 | A | SER | 59 | H |
| PRO | 373 | A | GLY | 63 | H |
| PHE | 374 | A | TYR | 58 | H |
| PHE | 375 | A | ASN | 34 | H |
| PHE | 375 | A | TYR | 37 | H |
| PHE | 375 | A | TYR | 58 | H |
| ASN | 437 | A | HIS | 57 | H |
| ASN | 437 | A | SER | 59 | H |
| ASN | 437 | A | ASN | 64 | H |
| ASN | 437 | A | TYR | 66 | H |
| SER | 438 | A | ASN | 64 | H |
| ASN | 439 | A | TYR | 38 | H |
| ASN | 439 | A | ASN | 64 | H |
| ASN | 439 | A | TYR | 66 | H |
| ASN | 439 | A | ASN | 114 | L |
| LYS | 440 | A | GLY | 63 | H |
| LYS | 440 | A | ASN | 64 | H |
| LYS | 440 | A | THR | 65 | H |
| LYS | 440 | A | TYR | 66 | H |
| LYS | 444 | A | ASN | 109 | L |
| PRO | 445 | A | ASN | 109 | L |
| PRO | 445 | A | SER | 113 | L |
| SER | 446 | A | ASN | 109 | L |
| ARG | 498 | A | LEU | 112 | H |
| ARG | 498 | A | ASP | 36 | L |
| ARG | 498 | A | ASN | 109 | L |
| PRO | 499 | A | TYR | 38 | H |
| PRO | 499 | A | TYR | 66 | H |
| PRO | 499 | A | VAL | 112 | H |
| PRO | 499 | A | ASN | 109 | L |
| PRO | 499 | A | SER | 113 | L |
| PRO | 499 | A | ASN | 114 | L |
| THR | 500 | A | VAL | 112 | H |
| THR | 500 | A | LEU | 112 | H |
| THR | 500 | A | ASP | 36 | L |
| THR | 500 | A | ASN | 37 | L |
| THR | 500 | A | TYR | 107 | L |
| THR | 500 | A | ASP | 108 | L |
| THR | 500 | A | ASN | 109 | L |
| THR | 500 | A | SER | 113 | L |
| THR | 500 | A | ASN | 114 | L |
| THR | 500 | A | VAL | 115 | L |
| TYR | 501 | A | VAL | 109 | H |
| TYR | 501 | A | GLY | 111 | H |
| TYR | 501 | A | VAL | 112 | H |
| TYR | 501 | A | LEU | 112 | H |
| GLY | 502 | A | VAL | 109 | H |
| GLY | 502 | A | GLY | 110 | H |
| GLY | 502 | A | GLY | 111 | H |
| GLY | 502 | A | VAL | 112 | H |
| GLY | 502 | A | LEU | 112 | H |
| VAL | 503 | A | TYR | 37 | H |
| VAL | 503 | A | HIS | 57 | H |
| VAL | 503 | A | VAL | 109 | H |
| VAL | 503 | A | GLY | 110 | H |
| VAL | 503 | A | GLY | 111 | H |
| GLY | 504 | A | VAL | 109 | H |
| GLY | 504 | A | GLY | 111 | H |
| HIS | 505 | A | GLY | 111 | H |
| GLN | 506 | A | TYR | 38 | H |
| GLN | 506 | A | HIS | 57 | H |
| GLN | 506 | A | ASN | 64 | H |
| GLN | 506 | A | TYR | 66 | H |
| GLN | 506 | A | VAL | 109 | H |
| GLN | 506 | A | VAL | 112 | H |
| GLN | 506 | A | ASN | 114 | L |
| TYR | 508 | A | HIS | 57 | H |

**Table S6. The list of the intermolecular contacts in the structure of the SA55 antibody complex with RBD (pdb id 7Y0W).** The interfacial contacts in the structure are defined by counting the number of interatomic contacts within a 5.5 Å distance threshold between atoms of the interacting proteins.

| **RBD Residue** | **RBD Residue Number** | **RBD chain** | **Ab Residue** | **Ab Residue Number** | **Ab chain** |
| --- | --- | --- | --- | --- | --- |
| PRO | 373 | R | LEU | 94 | B |
| PHE | 374 | R | THR | 57 | A |
| PHE | 374 | R | PHE | 55 | A |
| THR | 376 | R | PHE | 55 | A |
| ARG | 403 | R | PRO | 105 | A |
| ARG | 403 | R | ASN | 106 | A |
| GLY | 404 | R | PHE | 55 | A |
| GLY | 404 | R | LEU | 54 | A |
| GLY | 404 | R | ARG | 30 | A |
| ASP | 405 | R | LEU | 54 | A |
| ASP | 405 | R | SER | 31 | A |
| ASP | 405 | R | THR | 28 | A |
| ASP | 405 | R | ARG | 30 | A |
| GLU | 406 | R | ARG | 30 | A |
| VAL | 407 | R | ARG | 30 | A |
| VAL | 407 | R | PHE | 55 | A |
| VAL | 407 | R | LEU | 54 | A |
| ARG | 408 | R | ARG | 30 | A |
| ASN | 437 | R | ASP | 93 | B |
| ASN | 439 | R | TYR | 91 | B |
| ASN | 439 | R | ASP | 93 | B |
| LYS | 440 | R | ASP | 93 | B |
| VAL | 445 | R | HIS | 53 | B |
| TYR | 495 | R | PRO | 105 | A |
| SER | 496 | R | PRO | 105 | A |
| ARG | 498 | R | PHE | 112 | A |
| ARG | 498 | R | TYR | 49 | B |
| PRO | 499 | R | PHE | 100 | A |
| PRO | 499 | R | PRO | 101 | A |
| PRO | 499 | R | ASP | 50 | B |
| PRO | 499 | R | TYR | 91 | B |
| THR | 500 | R | GLY | 103 | A |
| THR | 500 | R | PHE | 112 | A |
| THR | 500 | R | PHE | 100 | A |
| THR | 500 | R | TYR | 49 | B |
| THR | 500 | R | ASP | 104 | A |
| THR | 500 | R | PRO | 101 | A |
| THR | 500 | R | ASN | 102 | A |
| THR | 500 | R | ASP | 50 | B |
| TYR | 501 | R | PRO | 105 | A |
| TYR | 501 | R | GLY | 103 | A |
| TYR | 501 | R | PHE | 112 | A |
| TYR | 501 | R | ASN | 102 | A |
| TYR | 501 | R | ASP | 104 | A |
| TYR | 501 | R | PRO | 101 | A |
| GLY | 502 | R | ASP | 104 | A |
| GLY | 502 | R | PRO | 101 | A |
| GLY | 502 | R | SER | 31 | A |
| GLY | 502 | R | GLY | 103 | A |
| GLY | 502 | R | HIS | 32 | A |
| GLY | 502 | R | ASN | 102 | A |
| VAL | 503 | R | PRO | 95 | B |
| VAL | 503 | R | PHE | 55 | A |
| VAL | 503 | R | VAL | 33 | A |
| VAL | 503 | R | ASN | 102 | A |
| VAL | 503 | R | LEU | 54 | A |
| VAL | 503 | R | HIS | 32 | A |
| VAL | 503 | R | PRO | 101 | A |
| VAL | 503 | R | ILE | 52 | A |
| VAL | 503 | R | SER | 31 | A |
| GLY | 504 | R | LEU | 54 | A |
| GLY | 504 | R | HIS | 32 | A |
| GLY | 504 | R | SER | 31 | A |
| GLY | 504 | R | ARG | 30 | A |
| HIS | 505 | R | PRO | 105 | A |
| HIS | 505 | R | HIS | 32 | A |
| HIS | 505 | R | ASP | 104 | A |
| HIS | 505 | R | SER | 31 | A |
| HIS | 505 | R | GLY | 103 | A |
| GLN | 506 | R | PRO | 101 | A |
| GLN | 506 | R | TYR | 91 | B |
| GLN | 506 | R | ASP | 93 | B |
| TYR | 508 | R | LEU | 54 | A |
| TYR | 508 | R | PHE | 55 | A |

**Table S7. The list of the intermolecular contacts in the structure of the XGI-171 antibody complex with RBD (pdb id 9KZD).** The interfacial contacts in the structure are defined by counting the number of interatomic contacts within a 5.5 Å distance threshold between atoms of the interacting proteins.

| **RBD Residue** | **RBD Residue Number** | **RBD chain** | **Ab Residue** | **Ab Residue Number** | **Ab chain** |
| --- | --- | --- | --- | --- | --- |
| TYR | 369 | A | ASP | 64 | H |
| TYR | 369 | A | TYR | 66 | H |
| PRO | 373 | A | LYS | 72 | H |
| PHE | 374 | A | LYS | 72 | H |
| PHE | 375 | A | LYS | 72 | H |
| PHE | 377 | A | TYR | 66 | H |
| PHE | 377 | A | PRO | 114 | L |
| LYS | 378 | A | ASP | 1 | L |
| LYS | 378 | A | SER | 109 | L |
| LYS | 378 | A | PRO | 114 | L |
| LYS | 378 | A | PRO | 115 | L |
| CYS | 379 | A | TYR | 66 | H |
| CYS | 379 | A | TYR | 108 | L |
| CYS | 379 | A | SER | 109 | L |
| CYS | 379 | A | MET | 113 | L |
| CYS | 379 | A | PRO | 114 | L |
| CYS | 379 | A | PRO | 115 | L |
| TYR | 380 | A | ILE | 2 | L |
| TYR | 380 | A | TYR | 108 | L |
| TYR | 380 | A | SER | 109 | L |
| TYR | 380 | A | MET | 113 | L |
| TYR | 380 | A | PRO | 114 | L |
| GLY | 381 | A | GLN | 111 | H |
| GLY | 381 | A | PHE | 38 | L |
| GLY | 381 | A | SER | 107 | L |
| GLY | 381 | A | TYR | 108 | L |
| GLY | 381 | A | SER | 109 | L |
| GLY | 381 | A | MET | 113 | L |
| VAL | 382 | A | GLN | 111 | H |
| VAL | 382 | A | TYR | 108 | L |
| VAL | 382 | A | MET | 113 | L |
| SER | 383 | A | GLY | 110 | H |
| SER | 383 | A | GLN | 111 | H |
| SER | 383 | A | PHE | 112 | H |
| SER | 383 | A | TRP | 113 | H |
| SER | 383 | A | MET | 113 | L |
| PRO | 384 | A | ASP | 64 | H |
| PRO | 384 | A | TYR | 66 | H |
| PRO | 384 | A | TRP | 113 | H |
| PRO | 384 | A | MET | 113 | L |
| PRO | 384 | A | PRO | 114 | L |
| THR | 385 | A | ASP | 38 | H |
| THR | 385 | A | ILE | 56 | H |
| THR | 385 | A | GLY | 57 | H |
| THR | 385 | A | THR | 58 | H |
| THR | 385 | A | ASP | 64 | H |
| THR | 385 | A | TYR | 66 | H |
| THR | 385 | A | TRP | 113 | H |
| LYS | 386 | A | ASP | 38 | H |
| LYS | 386 | A | GLY | 108 | H |
| LYS | 386 | A | SER | 109 | H |
| LYS | 386 | A | GLY | 110 | H |
| LYS | 386 | A | GLN | 111 | H |
| ASN | 388 | A | GLY | 59 | H |
| LEU | 390 | A | GLN | 111 | H |
| ALA | 411 | A | GLN | 27 | L |
| PRO | 412 | A | GLN | 27 | L |
| PRO | 412 | A | THR | 28 | L |
| PRO | 412 | A | TYR | 108 | L |
| GLY | 413 | A | GLN | 27 | L |
| PRO | 426 | A | TYR | 108 | L |
| ASP | 427 | A | THR | 28 | L |
| ASP | 427 | A | TYR | 108 | L |
| ASP | 428 | A | PHE | 38 | L |
| ASP | 428 | A | TYR | 108 | L |
| PHE | 429 | A | TYR | 108 | L |
| THR | 430 | A | TYR | 108 | L |

**Table S8. The list of the intermolecular contacts in the structure of the CR-3022 antibody complex with RBD (pdb id 6YM0).** The interfacial contacts in the structure are defined by counting the number of interatomic contacts within a 5.5 Å distance threshold between atoms of the interacting proteins.

| **RBD Residue** | **RBD Residue Number** | **RBD chain** | **Ab Residue** | **Ab Residue Number** | **Ab chain** |
| --- | --- | --- | --- | --- | --- |
| TYR | 369 | E | ILE | 30 | H |
| TYR | 369 | E | THR | 31 | H |
| TYR | 369 | E | GLY | 28 | H |
| TYR | 369 | E | PHE | 29 | H |
| TYR | 369 | E | TYR | 27 | H |
| ASN | 370 | E | GLY | 28 | H |
| ASN | 370 | E | TYR | 27 | H |
| SER | 371 | E | ILE | 30 | H |
| PHE | 374 | E | ILE | 30 | H |
| SER | 375 | E | TYR | 52 | H |
| SER | 375 | E | ILE | 30 | H |
| SER | 375 | E | GLY | 54 | H |
| THR | 376 | E | ILE | 30 | H |
| THR | 376 | E | TYR | 52 | H |
| THR | 376 | E | GLY | 54 | H |
| THR | 376 | E | ASP | 55 | H |
| PHE | 377 | E | THR | 31 | H |
| PHE | 377 | E | TRP | 33 | H |
| PHE | 377 | E | TYR | 52 | H |
| PHE | 377 | E | TYR | 32 | H |
| PHE | 377 | E | ILE | 30 | H |
| LYS | 378 | E | ILE | 30 | H |
| LYS | 378 | E | THR | 31 | H |
| LYS | 378 | E | TRP | 33 | H |
| LYS | 378 | E | ASP | 55 | H |
| LYS | 378 | E | GLU | 57 | H |
| LYS | 378 | E | TYR | 52 | H |
| CYS | 379 | E | THR | 31 | H |
| CYS | 379 | E | TRP | 33 | H |
| CYS | 379 | E | ILE | 102 | H |
| CYS | 379 | E | SER | 100 | H |
| CYS | 379 | E | GLY | 101 | H |
| TYR | 380 | E | TRP | 33 | H |
| TYR | 380 | E | ARG | 59 | H |
| TYR | 380 | E | THR | 104 | H |
| TYR | 380 | E | ILE | 102 | H |
| TYR | 380 | E | GLU | 57 | H |
| TYR | 380 | E | GLY | 101 | H |
| TYR | 380 | E | SER | 103 | H |
| GLY | 381 | E | ILE | 34 | L |
| GLY | 381 | E | TYR | 31 | L |
| GLY | 381 | E | THR | 104 | H |
| GLY | 381 | E | TYR | 38 | L |
| GLY | 381 | E | GLY | 101 | H |
| GLY | 381 | E | ILE | 102 | H |
| GLY | 381 | E | SER | 103 | H |
| GLY | 381 | E | TRP | 56 | L |
| VAL | 382 | E | GLY | 101 | H |
| VAL | 382 | E | ILE | 102 | H |
| VAL | 382 | E | SER | 103 | H |
| VAL | 382 | E | SER | 100 | H |
| VAL | 382 | E | TRP | 56 | L |
| VAL | 382 | E | ILE | 34 | L |
| VAL | 382 | E | THR | 104 | H |
| VAL | 382 | E | TYR | 38 | L |
| SER | 383 | E | GLY | 101 | H |
| SER | 383 | E | SER | 100 | H |
| SER | 383 | E | GLY | 99 | H |
| SER | 383 | E | THR | 104 | H |
| SER | 383 | E | PRO | 105 | H |
| PRO | 384 | E | SER | 100 | H |
| PRO | 384 | E | THR | 31 | H |
| PRO | 384 | E | GLY | 101 | H |
| THR | 385 | E | TYR | 32 | H |
| THR | 385 | E | THR | 31 | H |
| THR | 385 | E | SER | 100 | H |
| THR | 385 | E | GLN | 1 | H |
| THR | 385 | E | ASP | 107 | H |
| LYS | 386 | E | PRO | 105 | H |
| LYS | 386 | E | TYR | 55 | L |
| LYS | 386 | E | GLU | 61 | L |
| LYS | 386 | E | LEU | 52 | L |
| LYS | 386 | E | ASP | 107 | H |
| LYS | 386 | E | SER | 100 | H |
| ASP | 389 | E | TYR | 55 | L |
| LEU | 390 | E | TRP | 56 | L |
| PHE | 392 | E | TRP | 56 | L |
| PHE | 392 | E | ILE | 34 | L |
| ARG | 408 | E | ASP | 55 | H |
| ASP | 427 | E | TYR | 31 | L |
| ASP | 428 | E | SER | 32 | L |
| ASP | 428 | E | TYR | 31 | L |
| ASP | 428 | E | SER | 33 | L |
| ASP | 428 | E | TYR | 98 | L |
| PHE | 429 | E | TYR | 31 | L |
| THR | 430 | E | ILE | 34 | L |
| THR | 430 | E | TYR | 31 | L |
| THR | 430 | E | SER | 33 | L |
| THR | 430 | E | TYR | 38 | L |
| PHE | 515 | E | SER | 33 | L |
| PHE | 515 | E | ILE | 34 | L |
| GLU | 516 | E | ILE | 34 | L |
| GLU | 516 | E | SER | 33 | L |
| LEU | 517 | E | ILE | 34 | L |
| LEU | 517 | E | ASN | 35 | L |
| LEU | 517 | E | SER | 32 | L |
| LEU | 517 | E | SER | 33 | L |
| LEU | 517 | E | LYS | 36 | L |
| LEU | 518 | E | SER | 33 | L |
| HIS | 519 | E | ASN | 35 | L |

**Table S9. The list of the intermolecular contacts in the structure of the EY6A antibody complex with RBD (pdb id 7ZF3).** The interfacial contacts in the structure are defined by counting the number of interatomic contacts within a 5.5 Å distance threshold between atoms of the interacting proteins.

| **RBD Residue** | **RBD Residue Number** | **RBD chain** | **Ab Residue** | **Ab Residue Number** | **Ab chain** |
| --- | --- | --- | --- | --- | --- |
| LEU | 368 | E | TYR | 59 | H |
| LEU | 368 | E | ASN | 57 | H |
| TYR | 369 | E | ASN | 57 | H |
| TYR | 369 | E | LYS | 58 | H |
| TYR | 369 | E | SER | 56 | H |
| TYR | 369 | E | TYR | 59 | H |
| ASN | 370 | E | LYS | 58 | H |
| ASN | 370 | E | SER | 56 | H |
| ASN | 370 | E | TYR | 59 | H |
| ASN | 370 | E | ASN | 57 | H |
| ALA | 372 | E | LYS | 65 | H |
| PRO | 373 | E | GLY | 66 | H |
| PRO | 373 | E | LYS | 65 | H |
| PHE | 375 | E | LYS | 65 | H |
| THR | 376 | E | LYS | 65 | H |
| PHE | 377 | E | TYR | 59 | H |
| PHE | 377 | E | LYS | 65 | H |
| PHE | 377 | E | LEU | 95 | L |
| LYS | 378 | E | LEU | 95 | L |
| LYS | 378 | E | SER | 93 | L |
| LYS | 378 | E | ALA | 96 | L |
| LYS | 378 | E | ASP | 1 | L |
| LYS | 378 | E | ASP | 62 | H |
| CYS | 379 | E | LEU | 95 | L |
| CYS | 379 | E | SER | 93 | L |
| CYS | 379 | E | TYR | 92 | L |
| CYS | 379 | E | THR | 94 | L |
| TYR | 380 | E | SER | 93 | L |
| TYR | 380 | E | TYR | 92 | L |
| TYR | 380 | E | THR | 94 | L |
| GLY | 381 | E | SER | 93 | L |
| GLY | 381 | E | TYR | 92 | L |
| GLY | 381 | E | THR | 94 | L |
| GLY | 381 | E | TRP | 104 | H |
| GLY | 381 | E | SER | 91 | L |
| GLY | 381 | E | TYR | 32 | L |
| GLY | 381 | E | VAL | 105 | H |
| VAL | 382 | E | TRP | 104 | H |
| VAL | 382 | E | TYR | 32 | L |
| VAL | 382 | E | VAL | 105 | H |
| VAL | 382 | E | TYR | 92 | L |
| VAL | 382 | E | THR | 94 | L |
| SER | 383 | E | TRP | 104 | H |
| SER | 383 | E | VAL | 105 | H |
| SER | 383 | E | THR | 94 | L |
| SER | 383 | E | TYR | 106 | H |
| PRO | 384 | E | TYR | 59 | H |
| PRO | 384 | E | THR | 94 | L |
| PRO | 384 | E | TYR | 106 | H |
| PRO | 384 | E | LEU | 95 | L |
| PRO | 384 | E | ASN | 57 | H |
| THR | 385 | E | VAL | 50 | H |
| THR | 385 | E | TYR | 53 | H |
| THR | 385 | E | TYR | 106 | H |
| THR | 385 | E | SER | 52 | H |
| THR | 385 | E | TYR | 59 | H |
| THR | 385 | E | ASP | 33 | H |
| THR | 385 | E | ASN | 57 | H |
| THR | 385 | E | ILE | 51 | H |
| LYS | 386 | E | TRP | 104 | H |
| LYS | 386 | E | ASP | 33 | H |
| LYS | 386 | E | GLY | 101 | H |
| LYS | 386 | E | VAL | 105 | H |
| LYS | 386 | E | LEU | 103 | H |
| LYS | 386 | E | ASP | 99 | H |
| LYS | 386 | E | TYR | 106 | H |
| LYS | 386 | E | LYS | 102 | H |
| ASN | 388 | E | TYR | 53 | H |
| ASP | 389 | E | TYR | 53 | H |
| LEU | 390 | E | TRP | 104 | H |
| PHE | 392 | E | TRP | 104 | H |
| ALA | 411 | E | GLN | 27 | L |
| PRO | 412 | E | GLN | 27 | L |
| PRO | 412 | E | TYR | 92 | L |
| GLY | 413 | E | GLN | 27 | L |
| GLN | 414 | E | GLN | 27 | L |
| PRO | 426 | E | TYR | 92 | L |
| ASP | 427 | E | SER | 28 | L |
| ASP | 427 | E | SER | 30 | L |
| ASP | 427 | E | TYR | 92 | L |
| ASP | 428 | E | SER | 30 | L |
| ASP | 428 | E | TYR | 92 | L |
| PHE | 429 | E | TYR | 92 | L |
| THR | 430 | E | TYR | 92 | L |
| THR | 430 | E | TRP | 104 | H |
| LEU | 517 | E | TRP | 104 | H |
